# Efficient genetic perturbation of murine sensory neurons *in vivo* using CRISPR/Cas9

**DOI:** 10.1101/2025.10.20.683481

**Authors:** Guadalupe García, Jacob B. Shapiro, Zachary T. Campbell

## Abstract

Gene editing using CRISPR/Cas9 *in vivo* offers a powerful tool to investigate pain mechanisms. We validated the use of conditional knock-in mouse model expressing *Streptococcus pyogenes* CRISPR associated protein 9 selectively in sensory neurons by crossing with *Scn10a*-Cre driver. Transgene expression was confirmed in key tissues including the dorsal root ganglia (DRG) and sciatic nerve. To assess *in vivo* editing efficacy, RNA guides targeting GFP or TRPV1 were intrathecally administered. A dose of 3 µg RNA guides significantly reduced GFP expression, and two rounds of nanoparticle delivery targeting TRPV1 resulted in ∼65% reduction in DRG and ∼55% in sciatic nerve without triggering caspase-3-mediated apoptosis or motor deficits. Edited animals exhibited increased withdrawal latencies to heat and reduced nocifensive behaviors following capsaicin injection. Their responses to capsaicin-evoked thermal hyperalgesia and mechanical allodynia were diminished. This approach enables rapid and efficient sensory neuron-specific CRISPR/Cas9 gene perturbations for pain research in mice. We envisage that this method can be employed both for the exploration of molecular mechanisms underlying nociception and for the validation of therapeutic targets associated with pain.

## Introduction

Chronic pain is pervasive. It diminishes quality of life for more than 20% of the world’s population ^40^. In the vast majority of cases, pain is initiated by a specialized class of sensory neuron called a nociceptor. Their cell bodies reside in the dorsal root ganglia (DRG) and their fibers innervate internal organs, the viscera, and the skin. Nociceptors detect noxious thermal, mechanical, or chemical stimuli and transmit this information to the central nervous system. In chronic pain states, nociceptors become sensitized, leading to exaggerated responses to painful stimuli (hyperalgesia) and normally innocuous inputs (allodynia) ^17, 69^. Identifying genes that drive chronic pain through precise genetic changes is imperative for discovering analgesic targets ^5^. Although hundreds of genes have been linked to pain in mice ^22, 65, 77^ and humans ^46, 63, 72^, genetic validation remains extremely challenging due to the resource intensity of genome modification.

Genome engineering of rodents is a cornerstone of modern pain research. Numerous strategies including whole body mutation, conditional deletion, and temporally controlled knockouts, have enabled tremendous insights into pain signaling ^18, 39^. However, these approaches are time consuming, laborious, and expensive. A more rapid approach involves viral transduction. This strategy enables precise delivery of genetic cargo through intraganglionic injection of adeno-associated virus (AAV) vectors or retrograde transport of AAVs applied to the sciatic nerve ^61, 75, 76^. However, applying this approach in DRG presents significant challenges, including low and variable transduction efficiencies, transient expression, immune activation, and the limited packaging capacity of viral vectors — all of which restricts their utility ^39^.

CRISPR/Cas9 genome modification provides a precise and flexible approach to gene editing. SpCas9 nuclease is recruited to genomic targets via a programable non-coding RNA ^10, 16, 29^. When the guide-SpCas9 complex is bound to target DNA sequence, SpCas9 induces double-strand breaks that are repaired through the non-homologous end joining (NHEJ) pathway. This results in small insertions and deletions (indels) that often result in frameshifting and premature termination of translation. In the presence of a donor DNA template, homologous recombination (HR) can be used for genome engineering with desired replacements ^70^. Cas9 delivery via plasmid vectors or as a complex with preloaded guide RNAs has been reported in DRG sensory neurons ^44, 48, 60, 62^. The major goal of this work was to extend on this prior work by asking if highly efficient editing can be achieved through the use of stable transgenic expression of Cas9 in sensory neurons.

As a proof of concept, we targeted a well-characterized and widely studied ion channel critical to pain research: transient receptor potential vanilloid 1(TRPV1). TRPV1 is best known as the receptor of capsaicin, the compound responsible for the burning sensation of chili peppers and plays a central role in detecting noxious heat ^8^. TRPV1 has been implicated in both acute and chronic pain states ^6, 7, 51^. Employing guide RNAs to suppress TRPV1 expression in Cre-expressing Cas9 mice offers a precise and cell-specific approach to optimize genome engineering with behavioral and imaging-based endpoints. Here, we provide a proof of concept for rapid and facile genome perturbations within a subset of sensory neurons in mice. We envisage that our optimized strategy and genetic tool will be broadly applicable for the field.

## Materials and methods

### Mice

All animal procedures were conducted in accordance with approved protocols from the Institutional Animal Care and Use Committee (IACUC) at the University of Wisconsin-Madison (Protocol ID: M006582-A07). Rosa26-LSL-Cas9 knock-in (*Cas9^fl/+^*) mice were obtained from Jackson Laboratory (strain #024857). These mice express CRISPR-associated protein 9 (Cas9) endonuclease and EGFP in a Cre recombinase-dependent manner, under the control of a CAG promoter ^50^. Expression of Cas9 and EGFP is induced when a Cre gene is used. For this purpose, *Scn10a-Cre* (strain 036564) was used to generate conditional Cas9 knock-in mice ^59^. Then, *Cas9^fl/fl^/Scn10aCre^+^*, *Cas9^fl/fl^* and wild type C57BL/6J mice were obtained from our breeding facilities (Clinical Science Center, UW-Madison). All mice were housed under a 12-hour light/dark cycle with controlled ambient temperature (20–26°C) and humidity (30–70%). Food and water were provided *ad libitum*. Groups consisted of both male and female mice aged 8-12 weeks old (25-30 g) were used in this study. A cohort of mice was randomly assigned to either non-targeting control (NTC, 3 µg) or GFP (0.5-3 µg) RNA guide treatment, while a separate cohort was randomly assigned to NTC or TRPV1 RNA guide treatment. To minimize potential confounding factors, all animals were identified by their tag number and home cage. They were randomly assigned to experimental groups, ensuring that no group contained more than three mice from the same litter or the same cage. Cage positions within the rack were regularly rotated to avoid environmental or positional bias.

### Guide RNA design

The list of equipment needed for the production and validation of guide RNAs is provided in Table 1. The guide RNA synthesis protocol described here was based on that described by Dewitt et al. with minor modifications ^13^. For each target gene, three guides were designed against the gene of interest to ensure sufficient depletion (Fig. 1A). Candidate target (protospacer) sequences were identified with CHOPCHOP ^35^ using the GRCm38 genome assembly with the CRISPR/Cas9 and knockout settings. Protospacer sequences were selected on the basis of high predicted activity and specificity. The 20 nucleotide protospacer sequences were added to the T7FwdVar sequence (without the PAM) in place of the Xs in the T7FwdVar sequence (Table 2). The guanine upstream of the target sequence was required for efficient T7 transcription. If the target sequence begins with a guanine, the upstream guanine can be omitted.

**Figure 1.**
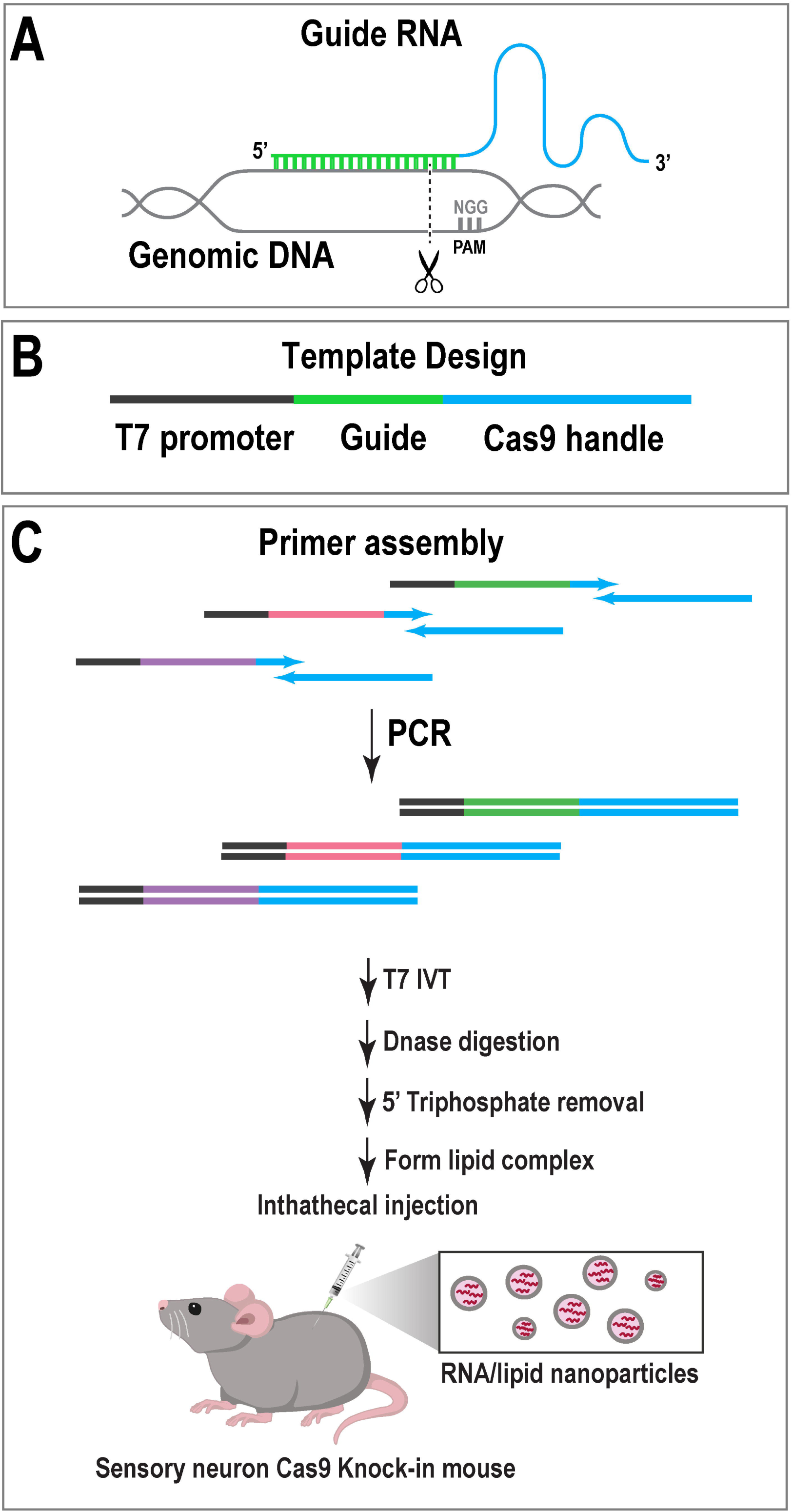
Overview of sgRNA synthesis and *in vivo* delivery. (**A**) Schematic of sgRNA binding to genomic DNA. (**B**) The DNA template for sgRNA synthesis comprises a T7 promoter, the gene-specific sgRNA sequence, and elements essential for Cas9 enzyme function. For each target gene, three distinct sgRNAs were designed to ensure sufficient gene depletion. (**C**) DNA templates were synthetized by PCR using overlapping primers. sgRNAs were produced with *In Vitro* transcription (IVT). Following DNase digestion of template and 5′-triphosphate removal, RNA-lipid complexes were prepared for intrathecal delivery into Cas9 knock-in mice.

**Table 1.** Key equipment, kits, and chemicals.

| Name | Manufacturer | Cat No |
| --- | --- | --- |
| Thermal cycler | Thermo Scientific | 4483636 |
| Tabletop centrifuge | Eppendorf | Centrifuge<br>5425R |
| Milli-Q Reference | EMD Millipore |  |
| 3Autoclave | Tuttnauer |  |
| Nanodrop One<sup>c</sup> | Thermo Scientific |  |
| RNA Clean & Concentrator-5 | Zymo Research | R1014 |
| GoTaq Green Master Mix | Promega | M7123 |
| HiScribe T7 High Yield RNA<br>Synthesis Kit | New England Biolabs | E2040L |
| DNase I, RNase-free | Thermo Scientific | EN0521 |
| Quick CIP | New England Biolabs | M0525S |
| Bleach | Clorox |  |
| RNase Away | Molecular Bioproducts | 7003 |
| DEPC | Sigma-Aldrich | 40718-100ML |
| Ethanol | Sigma-Aldrich | 1.08543.0250 |
| Agarose | Research Products International | A20090-500.0 |
| GeneRuler 1 kb Plus DNA<br>Ladder | Thermo Scientific | SM1331 |
| 50x TAE Buffer | Fisher Scientific | BP13321 |
| Ethidium Bromide | Fisher Scientific | BP102-5 |
| TriTrack DNA Loading Dye (6x) | Thermo Scientific | R1161 |
| DNA Oligonucleotides | IDT |  |

**Table 2:**
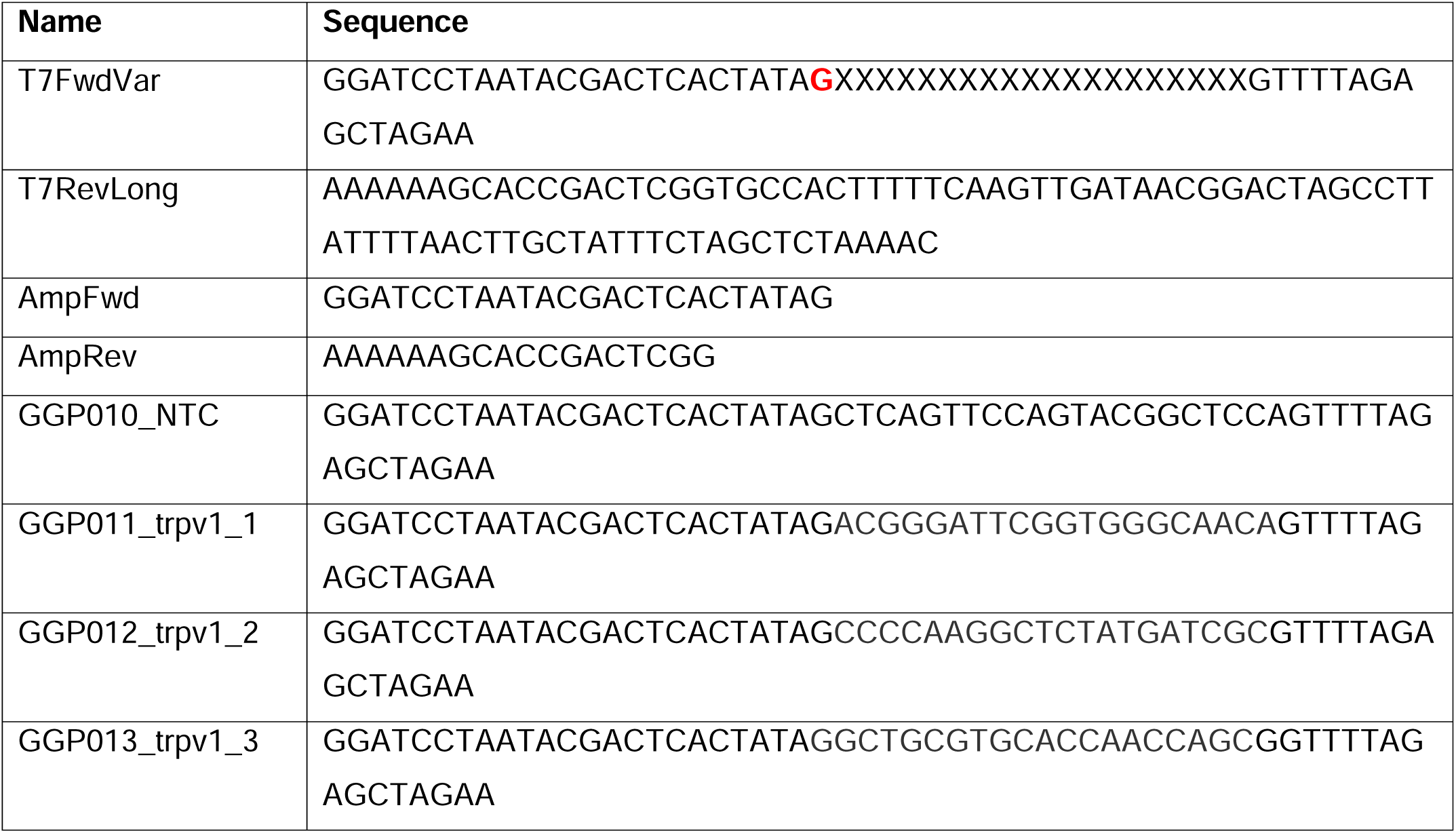
DNA oligonucleotides used in this study.

### Production of guide RNA pools

Prior to synthesis of guides, it is critical to mitigate the risk of contamination. RNase-free reagents, DEPC (diethyl pyrocarbonate)-treated water, and barrier pipette tips were used. To inactivate RNases with DEPC, 0.1% DEPC solution was prepared in ultrapure water and incubated overnight at 37°C. The solution was autoclaved with a sterilization time of at least 30 minutes. Benchtops and tube racks were decontaminated with 10% bleach for 15 minutes. The surfaces of electronic equipment were wiped down with RNase-away. Pipettes were disassembled and decontaminated with RNase-Away after soap, water, ethanol, and a bleach wash.

DNA templates for *in vitro* transcription (IVT) were created by PCR. The T7FwdVar and T7RevLong oligos served as the template for the full-length sgRNA (Table 2, Fig. 1B). FwdAmp and RevAmp prime amplification of the full-length template during subsequent rounds of PCR. GoTaq Green was added to a final volume of 25 μl for every individual guide (Table 3, Fig. 1C). The cycling parameters and extension times are provided in Table 4. The product was a 125 bp amplicon. To determine if a guide was amplified, products were visualized on a 2% TAE agarose gel (Fig. 2A).

**Figure 2.**
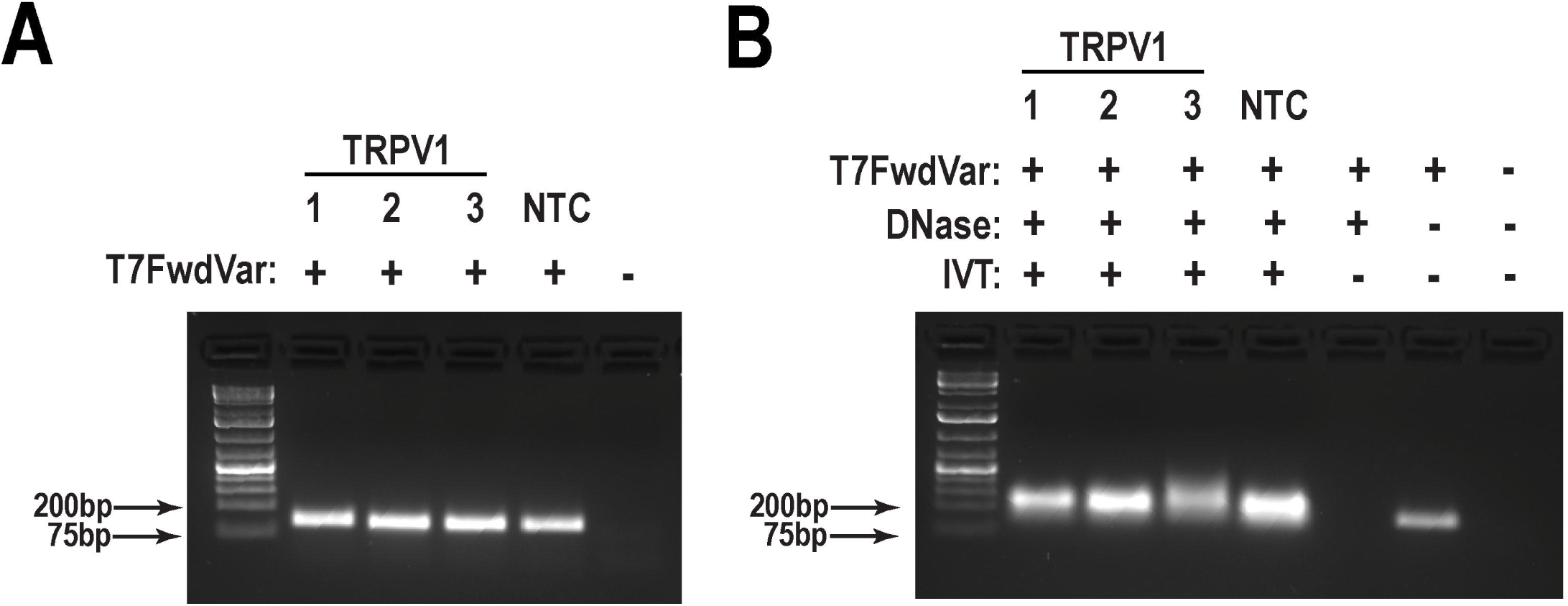
Validation of sgRNA synthesis (**A**) Successful amplification of the DNA templates (125 base pairs (bp)) for the targeted gene (TRPV1) or the non-targeting control (NTC) confirmed by agarose gel electrophoresis. (**B**) Successful *In vitro* transcription (IVT) of sgRNAs confirmed on a bleach-treated agarose gel. The presence of bands under ∼100-200 bp indicates successful sgRNA synthesis. These bands remained detectable following DNase treatment. No bands were observed in the absence of T7FwdVar primer or *in vitro* transcription.

**Table 3:** DNA template PCR conditions.

| Component | Concentration |
| --- | --- |
| GoTaq Green MM | 1x |
| T7FwdVarV2 | 40 nM |
| T7RevLongV2 | 40 nM |
| T7FwdAmp | 400 nM |
| T7RevAmp | 400 nM |

**Table 4:** sgRNA Template PCR Program.

| Temperature (C) | Duration | Cycles |
| --- | --- | --- |
| 98 | 1m | 1 |
| 98 | 15s | 30 |
| 50 | 15s |  |
| 72 | 15s |  |
| 72 | 2m | 1 |

Once suitable templates were created, the next step was *in vitro* transcription of RNA. Reagents from the HiScribe T7 High Yield RNA Synthesis kit – specifically the 10x reaction buffer and 100 mM nucleotide stocks were placed on ice for 1 hour. Next, a 2xIVT master mix was generated through the addition of equal volumes of 10x reaction buffer and 100 mM nucleotide stocks (Table 5). Individual reactions of 50 μl were prepared by mixing 23 μl of template PCR reaction, 23 μl of 2x IVT master mix, and 4 μl of T7 RNA Polymerase mix. Transcription was conducted at 37°C overnight.

**Table 5:** 2x In-vitro transcription master mix composition.

| Component | Concentration |
| --- | --- |
| T7 Reaction Buffer | 2x |
| ATP | 20 mM |
| CTP | 20 mM |
| GTP | 20 mM |
| UTP | 20 mM |

To degrade the DNA template, 2 U DNase I was added to each reaction and incubated for 15 minutes at 37°C. Next, RNA was purified using the Zymo RNA Clean and Concentrator-5 kit according to the manufacturer’s instructions with minor modifications. To reduce inflammation invoked by exogenous RNA carrying 5’-triphosphate groups, sgRNAs were incubated with calf intestinal alkaline phosphatase (Table 6, Fig. 1C) at 37°C for 3 hours. After the phosphatase reaction, sgRNAs were repurified with the Zymo RNA Clean and Concentrator kit, as described above. RNA was eluted in a total volume of 14 µl to concentrate the sample.

**Table 6:**
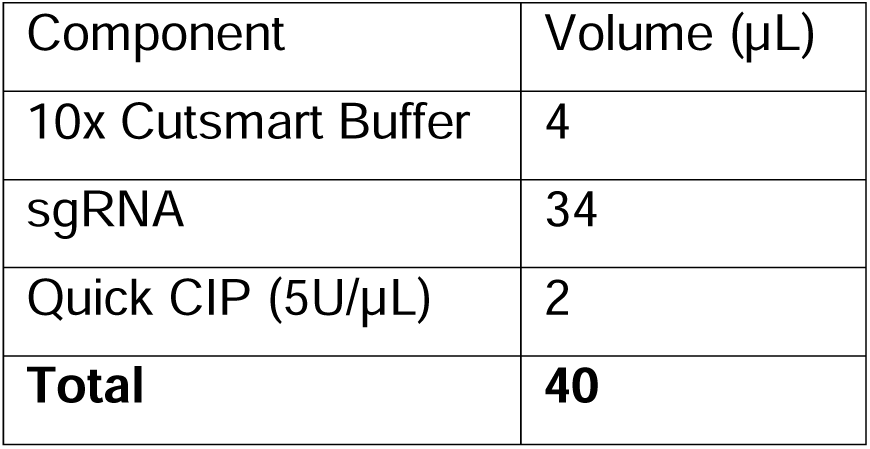
Quick CIP reaction conditions.

### Validation of guide RNAs

RNA quantification was conducted using a NanoDrop spectrophomoteter. This protocol produces highly concentrated RNA (∼5-10 μg/μL). To obtain accurate concentration measurements, samples were diluted 1:50 in DEPC-treated water before absorbance measurements were collected. To confirm the approximate size of the RNA, bleach gel electrophoresis was conducted ^2^. The gels are made with 1x TAE buffer to 2% (w/v) agarose and 1% (v/v) bleach. Bleach is allowed to react for 15 minutes prior to melting the agarose. Ethidium bromide at 0.5 μg/mL was added immediately prior to gel casting. A total of 100-500 ng of RNA was diluted in 10 μL in 1x TriTrack loading dye in DEPC-treated water and loaded onto the gel. Because these are not denaturing conditions, this gel simply confirms the presence of small (<200 nt) RNA molecules (Fig. 2B). More precise measurements can be collected on an Agilent TapeStation system if needed.

### Generation of guide RNA polymer complexes and intrathecal injection

After quality-control checks, individual sgRNAs targeting the same gene were pooled at equal concentrations. The final concentration of the sgRNA pool was 2-3 µg/µL. The remaining RNA was aliquoted and can be stored at −80°C for up to one month.

For intrathecal administration, the guide RNA was diluted with mRNA buffer to a final volume of 2 µL and mixed with 3 µL of *in vivo*-jetRNA^®^+ reagent (101000122, Polyplus). The resulting 5 µl of complex was loaded for injection using a Hamilton syringe equipped with a 31 G needle. A new needle was prepared for each mouse to avoid RNase contamination. Syringes were cleaned and sterilized between injections involving different guide RNA complexes. Mice were anesthetized using isoflurane at a concentration of 2–3% via a nose cone. Anesthesia depth was verified by the absence of a response to the tail or hind paw pinching. The lower lumbar region was shaved and disinfected with 70% ethanol. The injection site was identified between the L4–L5 vertebrae ^54^. The intrathecal injection was confirmed by a characteristic tail flick and the RNA complex volume was delivered slowly. Mice were returned to their home cages and monitored until full recovery from anesthesia, ensuring they were fully ambulatory and responsive. All procedures were conducted in a clean and decontaminated workspace.

### Immunohistochemistry

As editing targets, we examined either a transgenic GFP or endogenous TRPV1 using immunohistochemical analysis. Lumbar DRG, sciatic nerve, spinal cord or trigeminal ganglia were obtained from mice treated with guide RNAs targeting either: GFP (0.5-3 μg, a single intrathecal injection), TRPV1 or a non-targeting control (NTC) (3 µg, a single or double intrathecal injection). Mice were briefly anesthetized with isoflurane and transcardially perfused with ice-cold PBS followed by 4% formaldehyde (in PBS, pH=7.4). Tissues were harvested and post-fixed overnight at 4°C with 4% formaldehyde. Then, tissues were cryoprotected in 30% sucrose for 3 days at 4°C. Tissues were embedded in Tissue-Tek OCT compound (Sakura Finetek, Torrance, CA) and cryosectioned at 14 µm for DRG and trigeminal ganglia, and at 20 µm for sciatic nerve and spinal cord. Tissues sections were washed twice with TBS to remove OCT compound and then, incubated with 10 mM sodium citrate buffer (10 mM sodium citrate, 0.05% Tween 20, pH 6) for 1 hour. Tissues then were blocked with 10% normal goat serum, 0.3% Triton X-100 and goat anti-mouse Fab fragment (1:1,000, 115-001-003, Jackson ImmunoResearch) in TBS for 2 hours at room temperature using a humidified chamber. Sections were incubated overnight at 4°C with chicken anti-GFP (1:1,000, GFP-1010, Aves Labs), rabbit anti-GFP (1:200, 2956s, Cell Signaling Technology), mouse anti-Nav1.8 (1:1,000, 75-166, NeuroMab), chicken anti-peripherin (1:1,000, NBP1-05423, Novus Biologicals), rabbit anti-βIII-tubulin (1:1,000, 802001, Biolegends), goat anti-TRPV1 (1:500, GP14100, Neuromics), rabbit anti-caspase-3 (1:200, 9661, Cell Signaling Technology) or rabbit anti-CGRP (1:200, ab81887, Abcam) antibodies in 1% normal goat serum and 0.3% Triton X-100 as incubation solution. Sections were washed three times with 0.1%Triton X-100 in TBS and incubated with the corresponding secondary antibody (1:1,000, in incubation solution) or isolectin GS-IB4 (1:300, I21412, Invitrogen) for 1 hour at room temperature. Sections were washed and DAPI (1:10,000 in TBS) was incubated for 15 min followed by three more washes. Finally, coverslips were mounted on the slices using ProLong™ Glass Antifade (P36984, ThermoFisher Scientific).

### Confocal imaging and image analysis

Images were captured on a Leica SP8 laser scanning confocal microscope with LAS X Life Science software. Lumbar spinal cord sections were imaged using a 4x objective. Lumbar DRG, sciatic nerve and trigeminal ganglion sections were imaged using 20X objective or 2.5x zoom magnification. Imaging was performed with a pinhole of 1 Airy and scan speed of 400 Hz, using sequential acquisition to minimize spectral overlap. Images are representative projections of Z-stacks obtained from samples collected from 3-4 mice per experimental condition. Fluorescence intensity between treatments was quantified using ImageJ software by calculating corrected total cell fluorescence (CTCF) values. For DRG analysis, regions of interest (ROIs) were drawn to measure the TRPV1 or GFP fluoresce intensity specifically in GFP or Nav1.8 positive cells, respectively. For sciatic nerve, ROIs were created to measure the area of fibers in each fluorescence channel. Colocalization analysis was then performed to determine the area of overlap between markers in the sciatic nerve. For caspase-3 analysis, the number caspase-3-positive puncta was divided by the total number of GFP-positive cells and multiplied by 100 to calculate the percentage of positive cells. Results were calculated as either the mean of CTCF values or percent of at least 200 positive cells.

### Capsaicin-induced inflammatory pain model and behavioral tests

Intraplantar injection of capsaicin (5 μg/paw) elicits a rapid nocifensive response, characterized by flinching or licking of the injected paw. Therefore, we used capsaicin-induced inflammatory pain model to determine if delivery of TRPV1 RNA guides affects nocifensive behaviors ^3, 30, 57^. Mice were habituated to clear acrylic behavioral chambers for 1 hour per day over 2 days prior to the experimental days. Then, 20 μL of capsaicin stock solution (1 mg/mL, diluted in a solution containing 10% ethanol, 5% Tween-80 and 85% saline) were injected into the plantar surface of the left paw. The number of flinches was recorded during the first 10 minutes post-injection.

Forward looking infrared (FLIR) imaging was used to visualize thermal changes of the injected paw with capsaicin, using a FLIRT T31030sc thermal imaging camera (FLIR Systems, Wilsonville, OR) ^3, 47^. Mice were placed in acrylic boxes with wire mesh floors and allowed to acclimate 2 hours daily during the two days prior to testing, and for 30 min on the test day before the experiment. Mice were intraplantarly injected with 5 μg of capsaicin and placed into clear acrylic chambers boxes with wire mesh floor. The temperature of the medial plantar surface was registered twice, 1 h after capsaicin injection. The average temperature was then calculated and plotted.

To assess heat sensitivity, the hot plate test and Hargreaves test were employed ^12, 21^. Mice were habituated to the room for 1 hour daily on the two days prior to testing, and for 30 min before the start of the experiment. Then, mice were placed on a hot plate maintained at 50°C and the time taken to observe nocifensive behavior was recorded. Nocifensive behaviors were considered when mice showed hind paw withdrawal or licking, stamping, leaning posture, or jumping. The Hargreaves test was performed to measure paw withdrawal latency in response to a focused radiant heat source. Mice were placed on a glass floor within an enclosed chamber, and a high-intensity infrared light beam was directed at the plantar surface of the left hind paw. The light intensity was set at 25-30% of maximum output, as per the manufacturer’s guidelines (model 390, IITC). Paw withdrawal latency was recorded in triplicate, with a minimum interval of 10 minutes between trials to minimize experimental variability. The cutoff time was 20 seconds for both thermal tests. The average latency was then calculated and plotted.

Mechanical hypersensitivity was assessed using calibrated von Frey filaments (Stoelting, Wood Dale, IL, USA) to determine the paw withdrawal threshold in response to mechanical stimuli. Mice were placed in acrylic boxes with wire mesh floors and allowed to acclimate two hours daily during the two days prior to testing, and for one hour on the test day before the experiment. This procedure aimed to reduce stress-induced variability and obtain more homogeneous responses. Subsequently, von Frey filaments were applied to the plantar surface of the left hind paw for 2 seconds, and the up–down method ^9^ was performed to calculate the mechanical withdrawal threshold in grams (g) ^14^.

Motor coordination was assessed using rotarod test (Panlab, model LE8205) before and after i.t. injection of RNA guides. All the mice were habituated to the room conditions for at least 30 minutes before training. Mice previously received an adaptive training three consecutive days before the test started. The initial speed was 4 rpm and then, adjusted up to 10 rpm for 3 minutes. The time for each mouse to fall from the rod was recorded in each test as latency to fall.

During behavioral data acquisition, mice were randomly assigned to chambers for mechanical and thermal testing, or lanes for motor coordination assessment. At the end of each evaluation, the experimenter recorded the mouse’s tag number and home cage. The experimenter remained blinded to the RNA guides and treatments administered to the mice up to the final analysis.

### Western blotting

DRG, trigeminal ganglia (TG), spinal cord, thalamus and cerebral cortex were harvested from *Cas9^fl/fl^/Scn10aCre^+^*mice. Tissues were homogenized in ice-cold RIPA buffer supplemented with Pierce™ EDTA-free protease inhibitor (ThermoFisher, Waltham, MA, USA). Lysates were centrifugate at 4°C for 20 minutes at 12,000 *xg*. Protein concentration was quantified by Pierce™ BCA protein assay (Invitrogen, Waltham, MA, USA). Total protein (30 µg) was loaded onto an 8% SDS-polyacrylamide gel and subjected to gel electrophoresis. SDS-PAGE gel was then transferred to polyvinylidene difluoride membranes, followed by blocking with 5% non-fat dry milk in TBS plus 0.1% Tween-20 (TBS-T) for one hour. Membrane was then incubated overnight at 4°C in TBS-T with mouse anti-Cas9 (1:1,000; 14697, Cell Signaling Technology) or rabbit anti-GAPDH (1:20,000, 10494IAP, Proteintech). Membranes were then washed three times with 1% non-fat milk in TBS-T. Following washes, they were incubated with horseradish peroxidase (HRP)-conjugated anti-rabbit (1:10,000; 111-036-144, Jacskon Immunoresearch) or anti-mouse (1:5,000; 115-036-062, Jacskon Immunoresearch) secondary antibodies for one hour at room temperature. The protein signal was detected using the Immobilion ECL Ultra Western HRP substrate as chemiluminescence system (WBULS0100, Millipore) according to the manufactureŕs instructions. GAPDH was used as an internal loading control.

### Paclitaxel-induced cell death

To validate whether RNA guide treatment induced cell damage, we used paclitaxel-induced peripheral neuropathy as positive control of neurotoxicity and cellular cell damage ^23^. Mice were injected via intraperitoneal with paclitaxel (Cayman Chemical Company, 10461, Ann Arbor, MI) at dose of 4 mg/kg, administered every other day (a total of three injections). DRGs were obtained one day after last intraperitoneal injection.

### Statistical analysis of results

All data were analyzed using GraphPad Prism software. For comparisons between two groups, unpaired t-test was applied. For comparisons involving two variables, one or two-way ANOVA analysis followed by Dunnett or Bonferroni’s post hoc tests were used. Results were presented as mean ± SEM. Statistical significance was set at *P* < 0.05. Sample sizes were determined based on our prior work, which have consistently reported biologically and statistically meaningful results. Data were considered normally distributed if *P* > 0.05 and α= 0.05 by the Shapiro-Wilk test. For immunofluorescence analysis, tissue samples were obtained from 3 mice per condition. During sample collection, tissue processing, data acquisition, and quantitative analysis, all samples and data were recorded according to the mouse’s tag number. The experimenter remained blinded to the RNA guides and treatments administered to the mice until the final analysis. Data points were excluded if values were identified as outliers, defined as more than two standard deviations from the mean. The experimenter remained blinded to the genotype or RNA guides administered to the mice up to the final analysis.

For *in vivo* experiments, behavioral analyses were conducted using a sample size of 7 mice per group. Mice were included if they demonstrated consistent baselines responses to mechanical and thermal stimulation, as well as normal paw temperature and latency to fall, prior to the injection of RNA guides or drugs. After this point, no behavioral data were excluded from the analysis.

## Results

### Characterization of conditional Cas9 knock-in mice in sensory neurons

We generated a transgenic mouse model in which Cas9 is selectively expressed in sensory neurons by crossing Cas9*^fl/fl^* mice with mice expressing Cre recombinase under the control of the *Scn10a* promoter (Nav1.8-Cre) ^50, 59^. Since *Scn10a* encodes the voltage-gate sodium channel 1.8 (Nav1.8), which is primarily expressed in sensory neurons, we performed triple immunostaining to confirm that Cas9 expression is restricted to Nav1.8-positive neurons. We examined GFP as a reporter and co-labeled with peripherin marker which is co-expressed with Nav1.8 in over 90% of DRG sensory neurons (Fig. 3)^49, 52^. In conditional knock-in mice (*Cas9^fl/fl^/Scn10aCre^+^*), we found that neurons from lumbar DRGs (Fig. 3A) and trigeminal ganglia (Suppl. Fig.1A and 1B) exhibited GFP expression in Nav1.8 positive cells. In contrast, no GFP signal was observed in Cas9*^fl/fl^* mice lacking Cre expression, confirming Cre-dependent Cas9 activation. Furthermore, GFP-positive neurons colocalized with peripherin ^55^. GFP expression was also detected in sciatic nerve fibers (Fig. 3B), where it colocalized with Nav1.8 and βIII-tubulin, markers of neuronal fibers compared with samples under the absence of Cre recombinase. In addition, we found GFP signal specifically localized to laminae I and II of the spinal dorsal horn, where it colocalized with CGRP and IB4, markers of nociceptive primary inputs ^56, 78^. To demonstrate that the expression of Cas9 is restricted to sensory neurons, we performed immunodetection of Cas9 in pain pathway-related tissues obtained from conditional knock-in mice. We observed that Cas9 was expressed only in the DRG and TG, but not in the spinal cord, thalamus and cerebral cortex (Suppl. Fig.1C). These findings confirm that Cas9 expression is present in both DRG and TG sensory neurons in the *Cas9^fl/fl^/Scn10aCre^+^* transgenic model.

**Figure 3.**
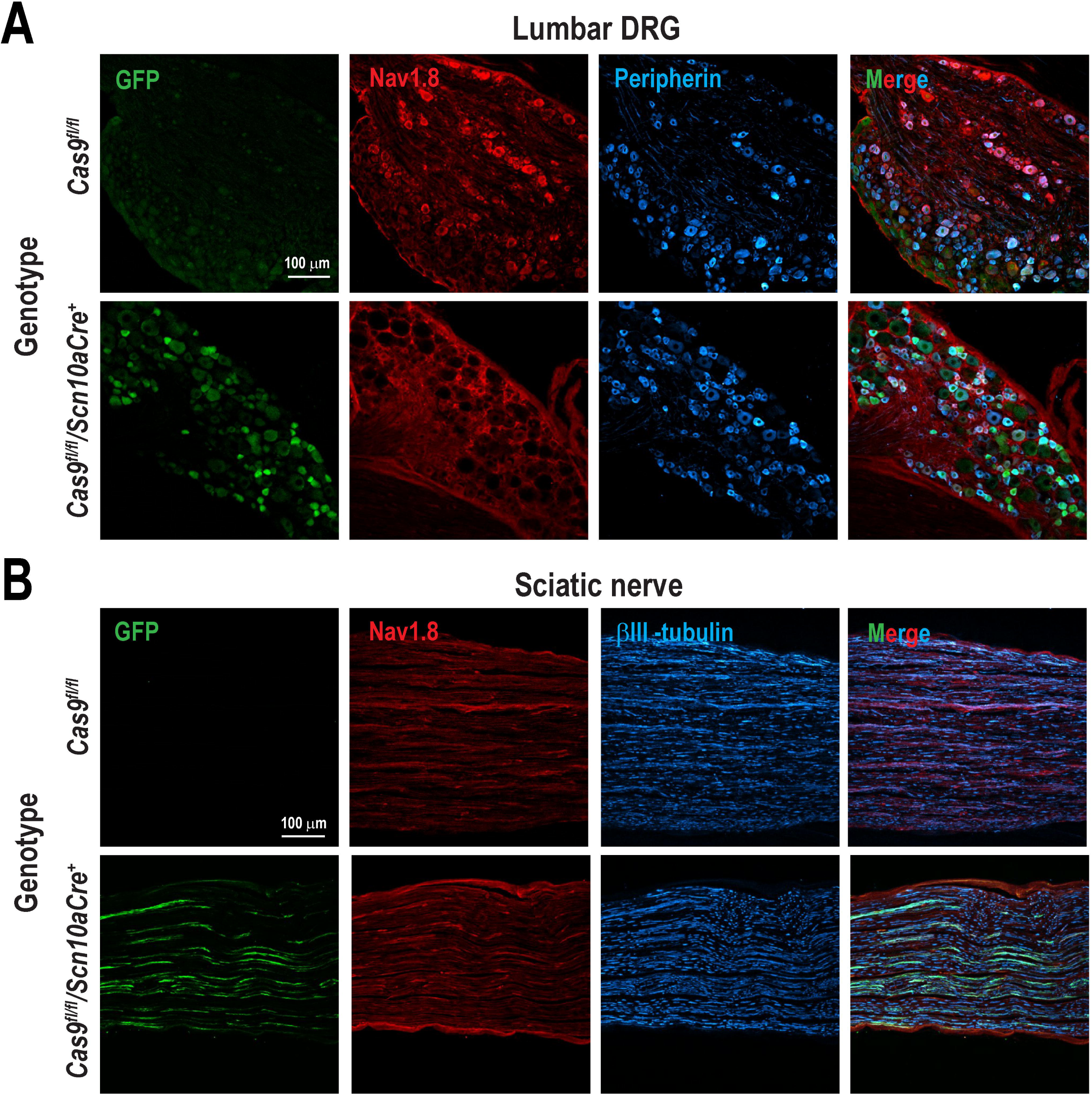
GFP expression in sensory neurons and fibers of Cas9 conditional knock-in mice. (**A**) Lumbar dorsal root ganglia (DRG) and (**B**) sciatic nerve sections from *Cas9^fl/fl^/Scn10aCre^+^*mice, in which expression of Cas9 endonuclease and GFP was induced in a Cre recombinase-dependent manner. No GFP expression was observed in absence of Cre recombinase (Cas9*^fl/fl^*). Cre recombinase was driven by the *Scn10a* (Nav1.8) promoter, selectively targeting sensory neurons. GFP signal indicated successful activation of the CRISPR/Cas9 system in Cre-expressing cells. Immunofluorescence analysis also showed colocalization of GFP with peripherin and βIII-tubulin, confirming selective Cas9 expression in small-diameter sensory neurons and neuronal fibers, respectively. Images are representative of three replicates.

### *In vivo* evaluation of RNA guide efficiency

Approximately 50% of guide RNAs are not effective at generating double-stranded breaks *in vivo* ^15, 53^. One challenge is the prediction of DNA accessibility. Additionally, in cell lines the use of more than three RNA guides simultaneously results in genotoxic stress ^1, 71^. To balance these two variables, we limited guide RNA pools to only 3, which should have an efficacy of 90% or more and minimal toxicity. As an initial approach to quantify editing efficacy, a range of guide RNA doses was injected that spanned 0.5, 1, 2 and 3 µg. The pools were either targeted to GFP or a non-targeting control (NTC) pool. We performed immunofluorescence analysis of DRG and sciatic nerve samples one week after RNA guide administration. We notably found that only the 3 µg dose significantly reduced GFP expression compared to the negative control group (Suppl. Fig. 2) (One-way ANOVA, F_(4,10)_ = 10.05, Dunnett’s test \*\**P* = 0.0015). Based on these findings, we selected the 3 µg dose for subsequent *in vivo* experiments.

To establish proof of concept, we focused on TRPV1 as a target. TRPV1 is an integral membrane protein that has been implicated in the transduction of noxious thermal stimuli and inflammatory pain ^7, 43, 64^. We evaluated the efficacy of RNA guides directed to diminish TRPV1 expression in DRG neurons. Injection of 3 μg RNA guide pools two times resulted in a significant decrease in TRPV1 signal of approximately 65% in the DRG (Fig. 4A) (Unpaired t-test, t_(4)_ = 3.27, \**P* = 0.03) and 55% in sciatic nerve (Fig. 4B) (Unpaired t-test, t_(4)_ = 3.88, \**P* = 0.02). These observations suggest that our strategy provides a robust means to achieve gene perturbation in murine DRG sensory neurons.

**Figure 4.**
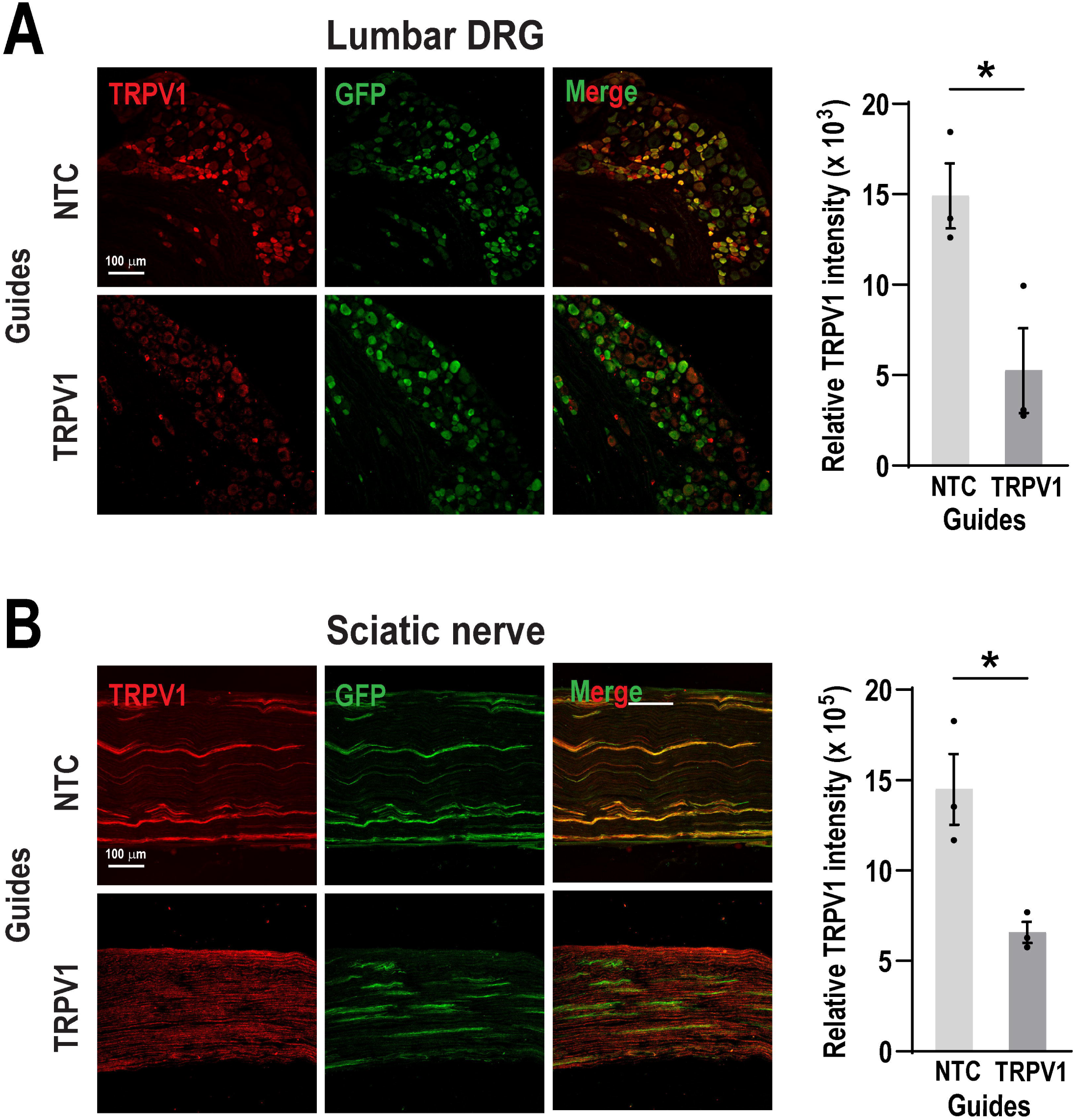
Intrathecal delivery of TRPV1 guides reduces TRPV1 expression *in vivo*. (**A**) DRG and (**B**) sciatic nerve sections showing immunohistochemical staining for TRPV1 expression following two intrathecal injections of 3 µg TRPV1 or NTC guides. TRPV1 fluorescence intensity was normalized to GFP signal. Quantification revealed a significant reduction in TRPV1 expression in TRPV1 guide-treated mice compared to NTC. Bars represent the mean ± SEM of relative intensity values from n = 3 mice per group. \**P* < 0.05 was determined by an unpaired t-test. Scale bar: 100 µm.

### Intrathecal TRPV1 guides does not induce caspase-3 activation

Caspase-3 is a critical effector of apoptosis, a form of programmed cell death. Upon activation, caspase-3 can degrade intracellular structural and functional proteins, ultimately leading to cell death ^28^. The CRISPR/Cas9 system induces double strand breaks in DNA, which can trigger DNA damage responses and, under certain conditions, initiate apoptotic signaling pathways ^20, 25^. To determine whether TRPV1 or NTC guide RNA pools induce DNA damage and apoptosis in DRG neurons, we assessed cleaved caspase-3 as a marker of apoptosis. We found that a minority of GFP positive cells showed cleaved caspase-3 puncta in either group, suggesting a low incidence of apoptosis (Unpaired t-test, t_(4)_ = 0.1, *P* = 0.92). As a positive control for apoptosis, we examined DRG cells from mice treated with paclitaxel (4 mg/kg, administered every other day for one week), a known inducer of neurotoxicity and cellular damage ^23^. Immunohistochemical analysis revealed that approximately 37% of DRG cells in paclitaxel-treated mice displayed immunoreactivity for caspase-3, confirming significant apoptotic activation in this condition compared with TRPV1 and NTC guide delivery (Fig. 5).

**Figure 5.**
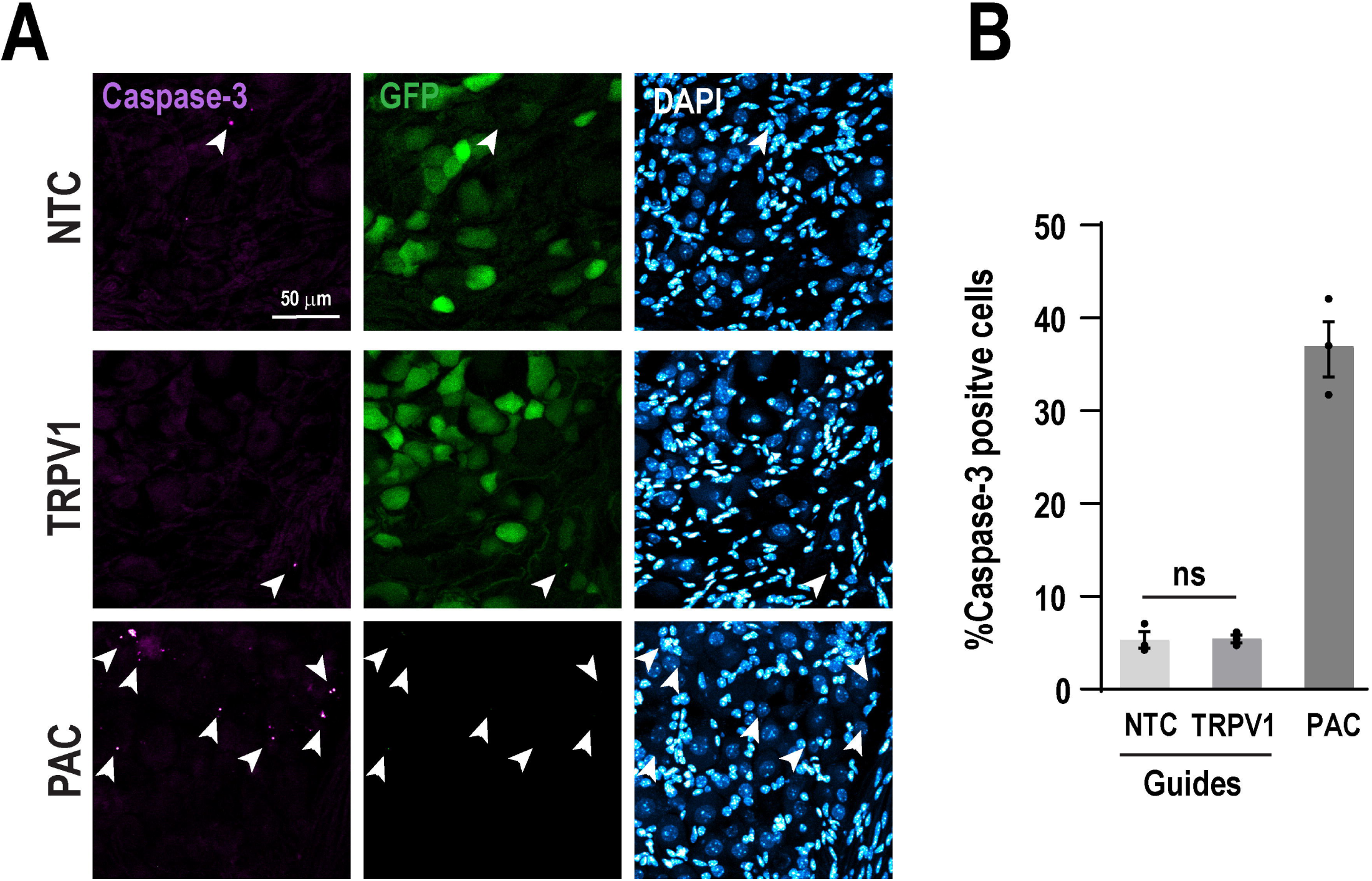
Perturbation of TRPV1 does not induce apoptosis. (**A**) Representative images of DRGs from mice treated with TRPV1 or NTC guides, stained for cleaved caspase-3 (apoptosis marker), GFP (Cas9 expression) and DAPI (nuclei). DRGs from mice treated with paclitaxel (4 mg/kg, administered every other day for one week) were included as a positive control of neurotoxicity and cellular damage. Scale bar: 50 µm. (**B**) Quantification of the percentage of cleaved caspase-3 positive cells among GFP positive neurons. Bars represent the mean ± SEM from n = 3 mice per group. No significant difference (ns) was determined by an unpaired t-test.

### TRPV1 guide delivery modulates heat-evoked behavioral responses

TRPV1 plays a crucial role in thermoregulation and is activated by temperatures above 42°C. It is also associated with the sensation of noxious heat and pain ^7, 8, 64^. We initially examined whether disruption of TRPV1 using conditional Cas9 knock-in mice with a single RNA guide delivery would affect behavioral heat responses. However, we observed that a single intrathecal guide injection failed to induce insensitivity to capsaicin-evoked thermal hyperalgesia and mechanical allodynia, as well as to reduce the increase in paw temperature in the injected side (Suppl. Fig. 3) (Two-way ANOVA, for thermal hyperalgesia: F_(4,50)_ = 0.37, *P* = 0.83; for mechanical allodynia: F_(8,75)_ = 1.36, *P* = 0.22; for paw temperature: unpaired t-test, t_(12)_ = 0.90, *P* = 0.38). We then administered a second intrathecal injection of the TRPV1 guide to determine whether capsaicin-evoked responses would be further affected. Our findings indicate that mice receiving two TRPV1 guide injections exhibited a delay in response latency of 11 seconds, compared to 6 seconds in the NTC group, when exposed to a 50°C heated plate (Fig. 6A) (Unpaired t-test, t_(12)_ = 5.02, \*\*\**P* < 0.001), Similarly, mice treated with TRPV1 guides showed increased response latency to radiant heat applied to the hind paw in the Hargreaves test, compared to the NTC group (Fig. 6B) (Unpaired t-test, t_(12)_ = 2.61, \**P* < 0.05). These findings suggest that intrathecal delivery of TRPV1 guides can modulate heat-evoked behavioral responses, likely through a reduction in TRPV1 expression in sensory neurons.

**Figure 6.**
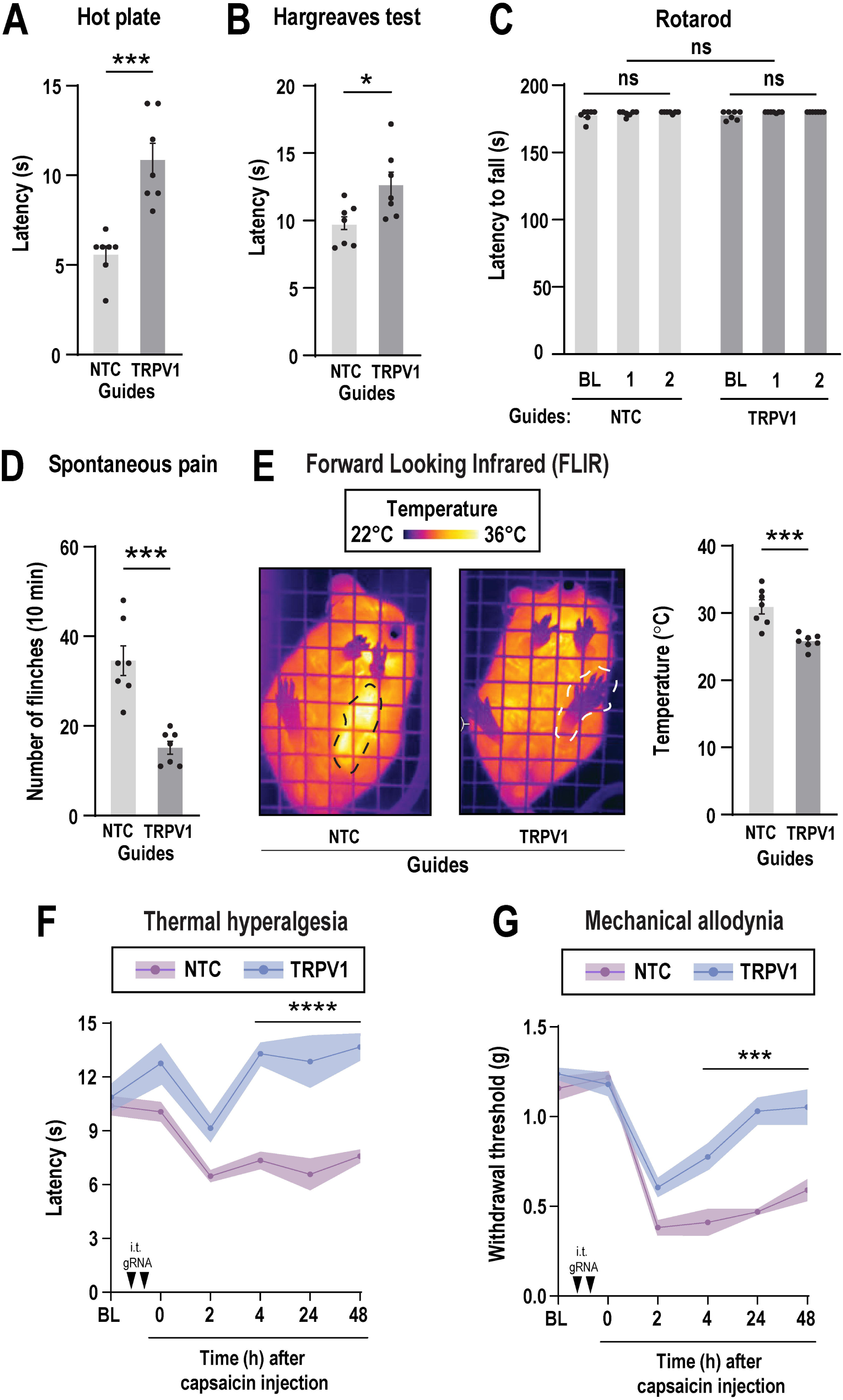
Perturbation of TRPV1 prevents capsaicin-induced hypersensitivity in Cas9 knock-in mice. Mice received two intrathecal injections of TRPV1 or not targeted control (NTC) guides (3 μg per injection) over a period of 2 weeks. One week after the last intrathecal injection, behavioral tests were conducted. Thermal hyperalgesia was assessed using (**A**) the hot plate test at 50°C and (**B**) the Hargreaves test. Bars represent mean ± SEM of withdrawal latency (s) from n = 7 mice per group. \**P* < 0.05 and \*\*\**P* < 0.001 was determined by an unpaired t-test. (**C**) Motor coordination was evaluated using the rotarod test at baseline (BL), and one week after the first (1) and second (2) intrathecal injection. Bars represent mean ± SEM of latency to fall (s) from n = 7 mice per group. No significant differences (ns) were observed across time points and RNA guide treatments by one-way repeated measures ANOVA. Capsaicin (5 μg) was intraplantarly injected and behavioral tests were conducted. (**D**) Spontaneous pain was assessed by counting the number of flinches during the first 10 minutes following intraplantar injection of capsaicin. Bars represent the mean ± SEM of number of flinches from n = 7 mice per group. \*\*\**P* < 0.001 was determined by an unpaired t-test. (**E**) Representative images of the left paw injected with capsaicin (dotted line) from TRPV1 or NTC guide-treated mice. Forward looking infrared (FLIR) imaging was used to measure paw temperature 1 h after capsaicin injection. Bars represent the mean ± SEM of temperature (°C) from n = 7 mice per group. \*\*\**P* < 0.001 was determined by an unpaired t-test. (**F**) Thermal hyperalgesia and (**G**) mechanical allodynia were evaluated before intrathecal injection of TRPV1 or NTC guides, as baseline (BL), and one week after second guide delivery at 0, 2, 4, 24 and 48 hours after capsaicin injection. Mice treated with TRPV1 guides displayed significantly reduced thermal and mechanical allodynia. Lines represent the mean ± a shaded area indicating SEM of withdrawal latency (s) or threshold (g) from n = 7 mice per group. \*\*\**P* < 0.001 and \*\*\*\**P* < 0.0001 were determined by two-way ANOVA followed by Bonferroni’s post hoc test. Arrows indicate intrathecal injections of RNA guides (i.t. gRNA).

To assess whether motor dysfunction could result from sgRNA delivery and DNA damage or intrathecal injection *per se*, we performed the rotarod test. Motor coordination and balance were evaluated by measuring the latency to fall from the rotating rod at baseline (prior to injection), and one week following the first and second intrathecal administration of TRPV1 or NTC guides. Our analysis revealed no significant differences in motor performance either within treated groups over time or between groups at any time point (Fig.7C) (Two-way ANOVA, F_(2.09, 12.59)_ = 1.76, *P* = 0.21). These results indicate that neither the delivery of sgRNAs nor the intrathecal injections adversely affected motor function.

### Intrathecal TRPV1 guides reduced capsaicin-induced pain behaviors

To determine if intrathecal delivery of TRPV1 guides impacts spontaneous pain associated behavior, we administered capsaicin (5 μg) into the paw. Capsaicin is a compound that selectively activates TRPV1 receptor ^34^. Intraplantar injection of capsaicin elicits a rapid nocifensive response, characterized by flinching or licking of the injected paw. We recorded this characteristic spontaneous pain behavior immediately after capsaicin injection over a 10-minute period. We observed that mice treated with TRPV1 guides displayed significantly fewer flinches than the NTC group (Fig. 6D) (Unpaired t-test, t_(12)_ = 5.42, \*\*\**P* < 0.001). As an additional test, we measured paw temperature using FLIR imaging. Heat is a cardinal hallmark of inflammation. We found capsaicin injection increased paw temperature on the ipsilateral (injected) paw in the NTC group but not in the TRPV1 edited group (Fig. 6E) (Unpaired t-test, t_(12)_ = 4.54, \*\*\**P* < 0.001), nor in the contralateral paw during the capsaicin test. (Suppl. Fig. 4) (Unpaired t-test, t_(12)_ = 0.75, *P* = 0.46). These results suggest that intrathecal delivery of TRPV1 guides can regulate pain and thermal responses evoked by capsaicin injection.

Next, we asked whether TRPV1 guides diminish sensitivity to a painful thermal challenge (hyperalgesia) and/or mechanical hypersensitivity to an innocuous stimulus (allodynia) following capsaicin injection. We found that capsaicin produced a reduction in thermal latency and mechanical withdrawal threshold in both the TRPV1 and NTC groups 2 h after injection. However, mice treated with TRPV1 guides recovered noticeably faster than mice that received NTC guides at 4, 24 and 48 h after capsaicin injection (Fig. 6F and 6G) (Two-way ANOVA, for thermal hyperalgesia: F_(5,60)_ = 4.98, Bonferroni’s multiple comparisons. \*\*\*\**P* < 0.0001; for mechanical allodynia: F_(5,60)_ = 6.74, Bonferroni’s multiple comparisons. \*\*\**P* < 0.001). The contralateral paw showed no changes in thermal or mechanical withdrawal responses throughout the capsaicin test (Suppl. Fig. 4) (Two-way ANOVA; for thermal hyperalgesia: F_(4,60)_ = 0.82, *P* = 0.51; for mechanical allodynia: F_(4,50)_ = 0.45, *P* = 0.77). These results suggest that intrathecal delivery of TRPV1 guides can regulate pain and thermal responses evoked by capsaicin injection. Based on these observations, we conclude that TRPV1 perturbation with CRISPR/Cas9 blunts capsaicin-induced evoked pain behaviors in mice.

## Discussion

Our approach leverages existing technologies and tools to achieve genetic perturbation in a subset of sensory neurons. We contextualize the major technical challenges and conceptual barriers with this strategy in the following section.

Editing efficiency is influenced by multiple factors. Cas9 dose influences editing rates when delivered by viruses ^67, 74^ or as a nanoparticle ^19^. We made use of transgenic expression as a means of holding Cas9 levels constant throughout adulthood. This perturbation of multiple genetic loci in a single animal, a challenging problem with standard approaches. While the *Scn10a* promoter offers the benefit of restricting Cas9 expression to a defined subset of sensory neurons, its expression is not strictly confined to nociceptors ^58^. While other strains such as *Advillin-Cre* and *Pirt-Cre* are used to drive Cre expression in sensory neurons, each has its own limitations regarding specificity ^24, 32^. Nevertheless, the general protocol we report here is likely comparable with other Cre lines that enable perturbation in different populations of sensory neurons.

A second variable is significant variation in the efficiency of guide RNAs. Approximately half of sgRNAs fail to induce double-strand breaks ^1, 53^. The use of guide pools improves editing efficiency by mitigating this risk ^36, 66, 73^. Anecdotally, we found poor editing efficiency in pilot experiments with single guide injections. We reasoned that making use of three guides should ensure success approximately 90% of the time. However, increasing the number of guides also raises the risk of off-target effects and cumulative DNA damage ^1, 71^. To balance efficacy and specificty, we limited each pool to three guides selected for high predicted activity and low off-target potential. A single 3 µg intrathecal dose of GFP guides reduced GFP expression *in vivo*, supporting the advantage of using multiple guides. This was further substantiated by editing of TRPV1, which achieved approximately 66% depletion after two rounds of guide RNA delivery.

A concern with the use of guide RNA pools is genotoxic stress. To determine whether the guide RNA pools we injected resulted in substantial DNA damage, we measured the expression of cleaved caspase-3. We did not observe a significant increase in apoptosis following guide RNA administration. Our approach is consistent with previous reports demonstrating that multi-guide delivery enhances editing efficiency without significantly increasing off-target risks through systemic delivery ^37, 66, 73^. Additionally, behavioral testing revealed no deficits in motor coordination, suggesting that our approach is well tolerated *in vivo*. Therefore, our approach aligns with current best practices for achieving efficient and specific editing in target tissues. A concern regarding the injection of exogenous RNAs is activation of the RIG-I pathway and induction of a type I interferon response. To mitigate this, we removed the 5’-triphosphate from our synthetic RNA guides ^68^. Our behavioral data suggest that the non-targeting guide RNA pools appeared to have minimal effects on behavior at baseline, further supporting the safety and tolerability of our approach.

We then applied this optimized strategy to target TRPV1, a key ion channel involved in noxious heat and capsaicin sensitivity across several pain models ^7, 8, 64^. While traditional TRPV1 knockout mice show impaired thermal nociception ^8, 42^, our initial results with delivery of a single TRPV1 guide failed to reproduce this phenotype. We found that mice remained sensitive to both thermal and mechanical capsaicin-evoked responses. This might be due to incomplete loss of TRPV1. Possible reasons for this include the inherent instability of RNA guides, which are rapidly degraded by endogenous RNases despite protective complex formation, limiting their bioavailability before reaching target tissues ^31^. Moreover, the efficiency of each guide is not uniform and may vary, even when computational predictions suggest high efficiency and specificity. To address this, we administered a second round of RNA delivery and validated knockdown efficacy through immunostaining of TRPV1. Following the second injection, we observed a significant reduction of TRPV1 expression in DRGs and sciatic nerve of approximately 65% and 55%, respectively. Similar results have been reported in human DRG neurons using CRISPR/Cas9 plasmid constructs, achieving a 57% reduction in TRPV1 protein levels ^48^. Our behavioral measurements are consistent with a reduction in TRPV1 function. Given these observations, it is likely that primary DRG sensory neurons after editing would display insensitivity to capsaicin *in vitro*. A limitation of this study is we did not probe if edited neurons displayed a reduction in TRPV1 function on sensory neuronal physiology.

How might this technology be leveraged for pain research? The major advantage of having stable Cas9 expression is the possibility of multiple rounds of genome editing in the same organism. This is potentially critical for understanding the function of paralogous genes. A notable example of this are voltage gated sodium channels. Considerable debate exists surrounding the necessity of Nav1.7 and Nav1.8 for pain ^26, 33^. Genetic deletion of Nav1.8 results in a loss of pain ^38^. The importance of Nav1.7 is more complicated. Embryonic loss of Nav1.7 in mouse or human sensory neurons results in profound analgesia ^11, 45^. Antagonists of either factor reduce pain in rodents ^4, 27, 41^. However, simultaneous deletion of Nav1.7 and Nav1.8 results in deficits in inflammatory pain but no effect on the development of neuropathic pain ^45^. This implies additional redundancy in voltage gated sodium channel function. One way to identify these factors is through the use of multi-locus genome perturbations. The strategy we report provides a means to this critical end but is only one example of a highly plastic tool for the field.

## ACKNOWLEDGEMENT

This work was supported by NIH grant R01NS114018 (Z.T.C.).

## AUTHOR CONTRIBUTIONS

J.B.S. was responsible for the molecular biology aspects of the manuscript. G.G. was responsible for the planning execution, analysis of all other aspects of the work and wrote the first draft of the manuscript. Z.T.C. contributed to the study conception assisted with the preparation of the manuscript and funding for the project.

## DECLARATION OF GENERATIVE AI AND AI-ASSISTED TECHNOLOGIES IN THE MANUSCRIPT PREPARATION PROCESS

During the preparation of this work the author(s) used ChatGPT (OpenAI) for editing to improve grammar, punctuation, and word choice. After using this tool, the author(s) reviewed and edited the content as needed and take(s) full responsibility for the content of the published article.

## DECLARATION OF INTERESTS

Z.T.C. is a consultant for Arrowhead Pharmaceuticals.

## References

1. Abdelrahman M, Wei Z, Rohila JS, Zhao K. Multiplex Genome-Editing Technologies for Revolutionizing Plant Biology and Crop Improvement. Front Plant Sci. 12:721203, 2021. doi:10.3389/fpls.2021.721203

2. Aranda PS, LaJoie DM, Jorcyk CL. Bleach gel: a simple agarose gel for analyzing RNA quality. Electrophoresis. 33:366–369, 2012. doi:10.1002/elps.201100335

3. Barragán-Iglesias P, Lou TF, Bhat VD, Megat S, Burton MD, Price TJ, Campbell ZT. Inhibition of Poly(A)-binding protein with a synthetic RNA mimic reduces pain sensitization in mice. Nat Commun. 9:10, 2018. doi:10.1038/s41467-017-02449-5

4. Beckley JT, Pajouhesh H, Luu G, Klas S, Delwig A, Monteleone D, Zhou X, Giuvelis D, Meng ID, Yeomans DC, Hunter JC, Mulcahy JV. Antinociceptive properties of an isoform-selective inhibitor of Nav1.7 derived from saxitoxin in mouse models of pain. Pain. 162:1250–1261, 2021. doi:10.1097/j.pain.0000000000002112

5. Berta T, Strong JA, Zhang JM, Ji RR. Targeting dorsal root ganglia and primary sensory neurons for the treatment of chronic pain: an update. Expert Opin Ther Targets. 27:665–678, 2023. doi:10.1080/14728222.2023.2247563

6. Bölcskei K, Helyes Z, Szabó Á, Sándor K, Elekes K, Németh J, Almási R, Pintér E, Pethő G, Szolcsányi J. Investigation of the role of TRPV1 receptors in acute and chronic nociceptive processes using gene-deficient mice. Pain. 117:368–376, 2005. doi:10.1016/j.pain.2005.06.024

7. Caterina MJ, Leffler A, Malmberg AB, Martin WJ, Trafton J, Petersen-Zeitz KR, Koltzenburg M, Basbaum AI, Julius D. Impaired nociception and pain sensation in mice lacking the capsaicin receptor. Science. 288:306–313, 2000. doi:10.1126/science.288.5464.306

8. Caterina MJ, Schumacher MA, Tominaga M, Rosen TA, Levine JD, Julius D. The capsaicin receptor: a heat-activated ion channel in the pain pathway. Nature. 389:816–824, 1997. doi:10.1038/39807

9. Chaplan SR, Bach FW, Pogrel JW, Chung JM, Yaksh TL. Quantitative assessment of tactile allodynia in the rat paw. J Neurosci Methods. 53:55–63, 1994. doi:10.1016/0165-0270(94)90144-9

10. Cho SW, Kim S, Kim JM, Kim JS. Targeted genome engineering in human cells with the Cas9 RNA-guided endonuclease. Nat Biotechnol. 31:230–232, 2013. doi:10.1038/nbt.2507

11. Cox JJ, Reimann F, Nicholas AK, Thornton G, Roberts E, Springell K, Karbani G, Jafri H, Mannan J, Raashid Y, Al-Gazali L, Hamamy H, Valente EM, Gorman S, Williams R, McHale DP, Wood JN, Gribble FM, Woods CG. An SCN9A channelopathy causes congenital inability to experience pain. Nature. 444:894–898, 2006. doi:10.1038/nature05413

12. Deuis JR, Dvorakova LS, Vetter I. Methods Used to Evaluate Pain Behaviors in Rodents. Front Mol Neurosci. 10:284, 2017. doi:10.3389/fnmol.2017.00284

13. Dewitt MW, J.; Wienert, B.; Schlapansky, M. F.; Aird, E.: In vitro transcription of guide RNAs and 5’-triphosphate removal V.12. Available at: https://www.protocols.io/view/in-vitro-transcription-of-guide-rnas-and-5-triphos-n2bvjyp5vk5w/v12?version_warning 2023

14. Dixon WJ. Efficient analysis of experimental observations. Annu Rev Pharmacol Toxicol. 20:441–462, 1980. doi:10.1146/annurev.pa.20.040180.002301

15. Doench JG, Hartenian E, Graham DB, Tothova Z, Hegde M, Smith I, Sullender M, Ebert BL, Xavier RJ, Root DE. Rational design of highly active sgRNAs for CRISPR-Cas9-mediated gene inactivation. Nat Biotechnol. 32:1262–1267, 2014. doi:10.1038/nbt.3026

16. Doudna JA, Charpentier E. Genome editing. The new frontier of genome engineering with CRISPR-Cas9. Science. 346:1258096, 2014. doi:10.1126/science.1258096

17. Dubin AE, Patapoutian A. Nociceptors: the sensors of the pain pathway. J Clin Invest. 120:3760–3772, 2010. doi:10.1172/jci42843

18. Eisener-Dorman AF, Lawrence DA, Bolivar VJ. Cautionary insights on knockout mouse studies: the gene or not the gene? Brain Behav Immun. 23:318–324, 2009. doi:10.1016/j.bbi.2008.09.001

19. Finn JD, Smith AR, Patel MC, Shaw L, Youniss MR, van Heteren J, Dirstine T, Ciullo C, Lescarbeau R, Seitzer J, Shah RR, Shah A, Ling D, Growe J, Pink M, Rohde E, Wood KM, Salomon WE, Harrington WF, Dombrowski C, Strapps WR, Chang Y, Morrissey DV. A Single Administration of CRISPR/Cas9 Lipid Nanoparticles Achieves Robust and Persistent In Vivo Genome Editing. Cell Reports. 22:2227–2235, 2018. doi:10.1016/j.celrep.2018.02.014

20. Haapaniemi E, Botla S, Persson J, Schmierer B, Taipale J. CRISPR–Cas9 genome editing induces a p53-mediated DNA damage response. Nature Medicine. 24:927–930, 2018. doi:10.1038/s41591-018-0049-z

21. Hargreaves K, Dubner R, Brown F, Flores C, Joris J. A new and sensitive method for measuring thermal nociception in cutaneous hyperalgesia. Pain. 32:77–88, 1988. 10.1016/0304-3959(88)90026-7

22. Hu G, Huang K, Hu Y, Du G, Xue Z, Zhu X, Fan G. Single-cell RNA-seq reveals distinct injury responses in different types of DRG sensory neurons. Sci Rep. 6:31851, 2016. doi:10.1038/srep31851

23. Huang ZZ, Li D, Liu CC, Cui Y, Zhu HQ, Zhang WW, Li YY, Xin WJ. CX3CL1-mediated macrophage activation contributed to paclitaxel-induced DRG neuronal apoptosis and painful peripheral neuropathy. Brain Behav Immun. 40:155–165, 2014. doi:10.1016/j.bbi.2014.03.014

24. Hunter DV, Smaila BD, Lopes DM, Takatoh J, Denk F, Ramer MS. Advillin Is Expressed in All Adult Neural Crest-Derived Neurons. eneuro. 5:ENEURO.0077-0018.2018, 2018. doi:10.1523/eneuro.0077-18.2018

25. Ihry RJ, Worringer KA, Salick MR, Frias E, Ho D, Theriault K, Kommineni S, Chen J, Sondey M, Ye C, Randhawa R, Kulkarni T, Yang Z, McAllister G, Russ C, Reece-Hoyes J, Forrester W, Hoffman GR, Dolmetsch R, Kaykas A. p53 inhibits CRISPR–Cas9 engineering in human pluripotent stem cells. Nature Medicine. 24:939–946, 2018. doi:10.1038/s41591-018-0050-6

26. Iseppon F, Kanellopoulos AH, Tian N, Zhou J, Caan G, Chiozzi R, Thalassinos K, Çubuk C, Lewis MJ, Cox JJ, Zhao J, Woods CG, Wood JN. Sodium channels Nav1.7, Nav1.8 and pain; two distinct mechanisms for Nav1.7 null analgesia. Neurobiology of Pain. 16:100168, 2024. 10.1016/j.ynpai.2024.100168

27. Jarvis MF, Honore P, Shieh CC, Chapman M, Joshi S, Zhang XF, Kort M, Carroll W, Marron B, Atkinson R, Thomas J, Liu D, Krambis M, Liu Y, McGaraughty S, Chu K, Roeloffs R, Zhong C, Mikusa JP, Hernandez G, Gauvin D, Wade C, Zhu C, Pai M, Scanio M, Shi L, Drizin I, Gregg R, Matulenko M, Hakeem A, Gross M, Johnson M, Marsh K, Wagoner PK, Sullivan JP, Faltynek CR, Krafte DS. A-803467, a potent and selective Nav1.8 sodium channel blocker, attenuates neuropathic and inflammatory pain in the rat. Proc Natl Acad Sci U S A. 104:8520–8525, 2007. doi:10.1073/pnas.0611364104

28. Jiang M, Qi L, Li L, Li Y. The caspase-3/GSDME signal pathway as a switch between apoptosis and pyroptosis in cancer. Cell Death Discovery. 6:112, 2020. doi:10.1038/s41420-020-00349-0

29. Jinek M, Chylinski K, Fonfara I, Hauer M, Doudna JA, Charpentier E. A programmable dual-RNA-guided DNA endonuclease in adaptive bacterial immunity. Science. 337:816–821, 2012. doi:10.1126/science.1225829

30. Joshi SK, Hernandez G, Mikusa JP, Zhu CZ, Zhong C, Salyers A, Wismer CT, Chandran P, Decker MW, Honore P. Comparison of antinociceptive actions of standard analgesics in attenuating capsaicin and nerve-injury-induced mechanical hypersensitivity. Neuroscience. 143:587–596, 2006. doi:10.1016/j.neuroscience.2006.08.005

31. Karikó K, Buckstein M, Ni H, Weissman D. Suppression of RNA Recognition by Toll-like Receptors: The Impact of Nucleoside Modification and the Evolutionary Origin of RNA. Immunity. 23:165–175, 2005. 10.1016/j.immuni.2005.06.008

32. Kim SH, Hadley SH, Maddison M, Patil M, Cha B, Kollarik M, Taylor-Clark TE. Mapping of Sensory Nerve Subsets within the Vagal Ganglia and the Brainstem Using Reporter Mice for Pirt, TRPV1, 5-HT3, and Tac1 Expression. eNeuro. 7, 2020. doi:10.1523/eneuro.0494-19.2020

33. King GF, Vetter I. No Gain, No Pain: NaV1.7 as an Analgesic Target. ACS Chemical Neuroscience. 5:749–751, 2014. doi:10.1021/cn500171p

34. Kwon DH, Zhang F, Suo Y, Bouvette J, Borgnia MJ, Lee S-Y. Heat-dependent opening of TRPV1 in the presence of capsaicin. Nature Structural & Molecular Biology. 28:554–563, 2021. doi:10.1038/s41594-021-00616-3

35. Labun K, Montague TG, Krause M, Torres Cleuren YN, Tjeldnes H, Valen E. CHOPCHOP v3: expanding the CRISPR web toolbox beyond genome editing. Nucleic Acids Res. 47:W171–w174, 2019. doi:10.1093/nar/gkz365

36. Li J, Kong D, Ke Y, Zeng W, Miki D. Application of multiple sgRNAs boosts efficiency of CRISPR/Cas9-mediated gene targeting in Arabidopsis. BMC Biology. 22:6, 2024. doi:10.1186/s12915-024-01810-7

37. Liang X, Potter J, Kumar S, Zou Y, Quintanilla R, Sridharan M, Carte J, Chen W, Roark N, Ranganathan S, Ravinder N, Chesnut JD. Rapid and highly efficient mammalian cell engineering via Cas9 protein transfection. Journal of Biotechnology. 208:44–53, 2015. 10.1016/j.jbiotec.2015.04.024

38. Lischka A, Lassuthova P, Çakar A, Record CJ, Van Lent J, Baets J, Dohrn MF, Senderek J, Lampert A, Bennett DL, Wood JN, Timmerman V, Hornemann T, Auer-Grumbach M, Parman Y, Hübner CA, Elbracht M, Eggermann K, Geoffrey Woods C, Cox JJ, Reilly MM, Kurth I. Genetic pain loss disorders. Nature Reviews Disease Primers. 8:41, 2022. doi:10.1038/s41572-022-00365-7

39. Luo L, Callaway EM, Svoboda K. Genetic Dissection of Neural Circuits: A Decade of Progress. Neuron. 98:256–281, 2018. doi:10.1016/j.neuron.2018.03.040

40. Lurie JM, Javaid A. Visualizing Global Chronic Pain. Anesth Analg. 138:918–919, 2024. doi:10.1213/ANE.0000000000006564

41. Minett MS, Falk S, Santana-Varela S, Bogdanov YD, Nassar MA, Heegaard AM, Wood JN. Pain without nociceptors? Nav1.7-independent pain mechanisms. Cell Rep. 6:301–312, 2014. doi:10.1016/j.celrep.2013.12.033

42. Mishra SK, Tisel SM, Orestes P, Bhangoo SK, Hoon MA. TRPV1-lineage neurons are required for thermal sensation. Embo j. 30:582–593, 2011. doi:10.1038/emboj.2010.325

43. Mitchell K, Lebovitz EE, Keller JM, Mannes AJ, Nemenov MI, Iadarola MJ. Nociception and inflammatory hyperalgesia evaluated in rodents using infrared laser stimulation after Trpv1 gene knockout or resiniferatoxin lesion. Pain. 155:733–745, 2014. doi:10.1016/j.pain.2014.01.007

44. Moreno AM, Alemán F, Catroli GF, Hunt M, Hu M, Dailamy A, Pla A, Woller SA, Palmer N, Parekh U, McDonald D, Roberts AJ, Goodwill V, Dryden I, Hevner RF, Delay L, Gonçalves Dos Santos G, Yaksh TL, Mali P. Long-lasting analgesia via targeted in situ repression of Na(V)1.7 in mice. Sci Transl Med. 13, 2021. doi:10.1126/scitranslmed.aay9056

45. Nassar MA, Stirling LC, Forlani G, Baker MD, Matthews EA, Dickenson AH, Wood JN. Nociceptor-specific gene deletion reveals a major role for Nav1.7 (PN1) in acute and inflammatory pain. Proc Natl Acad Sci U S A. 101:12706–12711, 2004. doi:10.1073/pnas.0404915101

46. Nguyen MQ, von Buchholtz LJ, Reker AN, Ryba NJ, Davidson S. Single-nucleus transcriptomic analysis of human dorsal root ganglion neurons. Elife. 10, 2021. doi:10.7554/eLife.71752

47. Nosrati Z, Bergamo M, Rodríguez-Rodríguez C, Saatchi K, Häfeli UO. Refinement and validation of infrared thermal imaging (IRT): a non-invasive technique to measure disease activity in a mouse model of rheumatoid arthritis. Arthritis Res Ther. 22:281, 2020. doi:10.1186/s13075-020-02367-w

48. Palomino SM, Gabriel KA, Mwirigi JM, Cervantes A, Horton P, Funk G, Moutal A, Martin LF, Khanna R, Price TJ, Patwardhan A. Genetic editing of primary human dorsal root ganglion neurons using CRISPR-Cas9. Sci Rep. 15:11116, 2025. doi:10.1038/s41598-025-91153-2

49. Patil MJ, Hovhannisyan AH, Akopian AN. Characteristics of sensory neuronal groups in CGRP-cre-ER reporter mice: Comparison to Nav1.8-cre, TRPV1-cre and TRPV1-GFP mouse lines. PLOS ONE. 13:e0198601, 2018. doi:10.1371/journal.pone.0198601

50. Platt RJ, Chen S, Zhou Y, Yim MJ, Swiech L, Kempton HR, Dahlman JE, Parnas O, Eisenhaure TM, Jovanovic M, Graham DB, Jhunjhunwala S, Heidenreich M, Xavier RJ, Langer R, Anderson DG, Hacohen N, Regev A, Feng G, Sharp PA, Zhang F. CRISPR-Cas9 knockin mice for genome editing and cancer modeling. Cell. 159:440–455, 2014. doi:10.1016/j.cell.2014.09.014

51. Rahman MM, Jo YY, Kim YH, Park CK. Current insights and therapeutic strategies for targeting TRPV1 in neuropathic pain management. Life Sci. 355:122954, 2024. doi:10.1016/j.lfs.2024.122954

52. Ramachandra R, Elmslie KS. Voltage-dependent sodium (NaV) channels in group IV sensory afferents. Molecular Pain. 12:1744806916660721, 2016. doi:10.1177/1744806916660721

53. Ran FA, Cong L, Yan WX, Scott DA, Gootenberg JS, Kriz AJ, Zetsche B, Shalem O, Wu X, Makarova KS, Koonin EV, Sharp PA, Zhang F. In vivo genome editing using Staphylococcus aureus Cas9. Nature. 520:186–191, 2015. doi:10.1038/nature14299

54. Rigaud M, Gemes G, Barabas ME, Chernoff DI, Abram SE, Stucky CL, Hogan QH. Species and strain differences in rodent sciatic nerve anatomy: implications for studies of neuropathic pain. Pain. 136:188–201, 2008. doi:10.1016/j.pain.2008.01.016

55. Romano R, Del Fiore VS, Bucci C. Role of the Intermediate Filament Protein Peripherin in Health and Disease. Int J Mol Sci. 23, 2022. doi:10.3390/ijms232315416

56. Saeed AW, Ribeiro-da-Silva A. Non-peptidergic primary afferents are presynaptic to neurokinin-1 receptor immunoreactive lamina I projection neurons in rat spinal cord. Mol Pain. 8:64, 2012. doi:10.1186/1744-8069-8-64

57. Sawynok J, Reid A, Meisner J. Pain behaviors produced by capsaicin: influence of inflammatory mediators and nerve injury. J Pain. 7:134–141, 2006. doi:10.1016/j.jpain.2005.09.013

58. Shields SD, Ahn H-S, Yang Y, Han C, Seal RP, Wood JN, Waxman SG, Dib-Hajj SD. Nav1.8 expression is not restricted to nociceptors in mouse peripheral nervous system. PAIN®. 153:2017–2030, 2012. 10.1016/j.pain.2012.04.022

59. Stirling LC, Forlani G, Baker MD, Wood JN, Matthews EA, Dickenson AH, Nassar MA. Nociceptor-specific gene deletion using heterozygous NaV1.8-Cre recombinase mice. Pain. 113:27–36, 2005. doi:10.1016/j.pain.2004.08.015

60. Stratton HJ, Dolatyari M, Kopruszinski C, Ghetti A, Maciuba S, Bowden G, Rivière P, Barber K, Dodick DW, Edorh E, Dumaire N, Moutal A, Navratilova E, Porreca F. A prolactin-targeting antibody to prevent stress-induced peripheral nociceptor sensitization and female postoperative pain. Proc Natl Acad Sci U S A. 122:e2501229122, 2025. doi:10.1073/pnas.2501229122

61. Tadokoro T, Bravo-Hernandez M, Agashkov K, Kobayashi Y, Platoshyn O, Navarro M, Marsala S, Miyanohara A, Yoshizumi T, Shigyo M, Krotov V, Juhas S, Juhasova J, Nguyen D, Kupcova Skalnikova H, Motlik J, Studenovska H, Proks V, Reddy R, Driscoll SP, Glenn TD, Kemthong T, Malaivijitnond S, Tomori Z, Vanicky I, Kakinohana M, Pfaff SL, Ciacci J, Belan P, Marsala M. Precision spinal gene delivery-induced functional switch in nociceptive neurons reverses neuropathic pain. Mol Ther. 30:2722–2745, 2022. doi:10.1016/j.ymthe.2022.04.023

62. Tao Y, Wang QH, Li XT, Liu Y, Sun RH, Xu HJ, Zhang M, Li SY, Yang L, Wang HJ, Hao LY, Cao JL, Pan Z. Spinal-Specific Super Enhancer in Neuropathic Pain. J Neurosci. 43:8547–8561, 2023. doi:10.1523/jneurosci.1006-23.2023

63. Tavares-Ferreira D, Shiers S, Ray PR, Wangzhou A, Jeevakumar V, Sankaranarayanan I, Cervantes AM, Reese JC, Chamessian A, Copits BA, Dougherty PM, Gereau RWt, Burton MD, Dussor G, Price TJ. Spatial transcriptomics of dorsal root ganglia identifies molecular signatures of human nociceptors. Sci Transl Med. 14:eabj8186, 2022. doi:10.1126/scitranslmed.abj8186

64. Tominaga M, Caterina MJ, Malmberg AB, Rosen TA, Gilbert H, Skinner K, Raumann BE, Basbaum AI, Julius D. The cloned capsaicin receptor integrates multiple pain-producing stimuli. Neuron. 21:531–543, 1998. doi:10.1016/s0896-6273(00)80564-4

65. Usoskin D, Furlan A, Islam S, Abdo H, Lönnerberg P, Lou D, Hjerling-Leffler J, Haeggström J, Kharchenko O, Kharchenko PV, Linnarsson S, Ernfors P. Unbiased classification of sensory neuron types by large-scale single-cell RNA sequencing. Nature Neuroscience. 18:145–153, 2015. doi:10.1038/nn.3881

66. Wang H, Yang H, Shivalila CS, Dawlaty MM, Cheng AW, Zhang F, Jaenisch R. One-Step Generation of Mice Carrying Mutations in Multiple Genes by CRISPR/Cas-Mediated Genome Engineering. Cell. 153:910–918, 2013. 10.1016/j.cell.2013.04.025

67. Wang L, Yang Y, Breton CA, White J, Zhang J, Che Y, Saveliev A, McMenamin D, He Z, Latshaw C, Li M, Wilson JM. CRISPR/Cas9-mediated in vivo gene targeting corrects hemostasis in newborn and adult factor IX–knockout mice. Blood. 133:2745–2752, 2019. doi:10.1182/blood.2019000790

68. Wienert B, Shin J, Zelin E, Pestal K, Corn JE. In vitro–transcribed guide RNAs trigger an innate immune response via the RIG-I pathway. PLOS Biology. 16:e2005840, 2018. doi:10.1371/journal.pbio.2005840

69. Woolf CJ, Ma Q. Nociceptors--noxious stimulus detectors. Neuron. 55:353–364, 2007. doi:10.1016/j.neuron.2007.07.016

70. Xue C, Greene EC. DNA Repair Pathway Choices in CRISPR-Cas9-Mediated Genome Editing. Trends Genet. 37:639–656, 2021. doi:10.1016/j.tig.2021.02.008

71. Yang L, Li H, Han Y, Song Y, Wei M, Fang M, Sun Y. CRISPR/Cas9 Gene Editing System Can Alter Gene Expression and Induce DNA Damage Accumulation. Genes (Basel). 14, 2023. doi:10.3390/genes14040806

72. Yang L, Xu M, Bhuiyan SA, Li J, Zhao J, Cohrs RJ, Susterich JT, Signorelli S, Green U, Stone JR, Levy D, Lennerz JK, Renthal W. Human and mouse trigeminal ganglia cell atlas implicates multiple cell types in migraine. Neuron. 110:1806–1821.e1808, 2022. doi:10.1016/j.neuron.2022.03.003

73. Yin C, Zhang T, Qu X, Zhang Y, Putatunda R, Xiao X, Li F, Xiao W, Zhao H, Dai S, Qin X, Mo X, Young WB, Khalili K, Hu W. In Vivo Excision of HIV-1 Provirus by saCas9 and Multiplex Single-Guide RNAs in Animal Models. Mol Ther. 25:1168–1186, 2017. doi:10.1016/j.ymthe.2017.03.012

74. Yin H, Song CQ, Dorkin JR, Zhu LJ, Li Y, Wu Q, Park A, Yang J, Suresh S, Bizhanova A, Gupta A, Bolukbasi MF, Walsh S, Bogorad RL, Gao G, Weng Z, Dong Y, Koteliansky V, Wolfe SA, Langer R, Xue W, Anderson DG. Therapeutic genome editing by combined viral and non-viral delivery of CRISPR system components in vivo. Nat Biotechnol. 34:328–333, 2016. doi:10.1038/nbt.3471

75. Yu H, Fischer G, Hogan QH. AAV-Mediated Gene Transfer to Dorsal Root Ganglion. Methods Mol Biol. 1382:251–261, 2016. doi:10.1007/978-1-4939-3271-9_18

76. Zheng CX, Wang SM, Bai YH, Luo TT, Wang JQ, Dai CQ, Guo BL, Luo SC, Wang DH, Yang YL, Wang YY. Lentiviral Vectors and Adeno-Associated Virus Vectors: Useful Tools for Gene Transfer in Pain Research. Anat Rec (Hoboken). 301:825–836, 2018. doi:10.1002/ar.23723

77. Zheng Y, Liu P, Bai L, Trimmer JS, Bean BP, Ginty DD. Deep Sequencing of Somatosensory Neurons Reveals Molecular Determinants of Intrinsic Physiological Properties. Neuron. 103:598–616.e597, 2019. doi:10.1016/j.neuron.2019.05.039

78. Zylka MJ, Rice FL, Anderson DJ. Topographically distinct epidermal nociceptive circuits revealed by axonal tracers targeted to Mrgprd. Neuron. 45:17–25, 2005. doi:10.1016/j.neuron.2004.12.015

