## Supplement for "Efficient genetic perturbation of murine sensory neurons *in vivo* using CRISPR/Cas9"

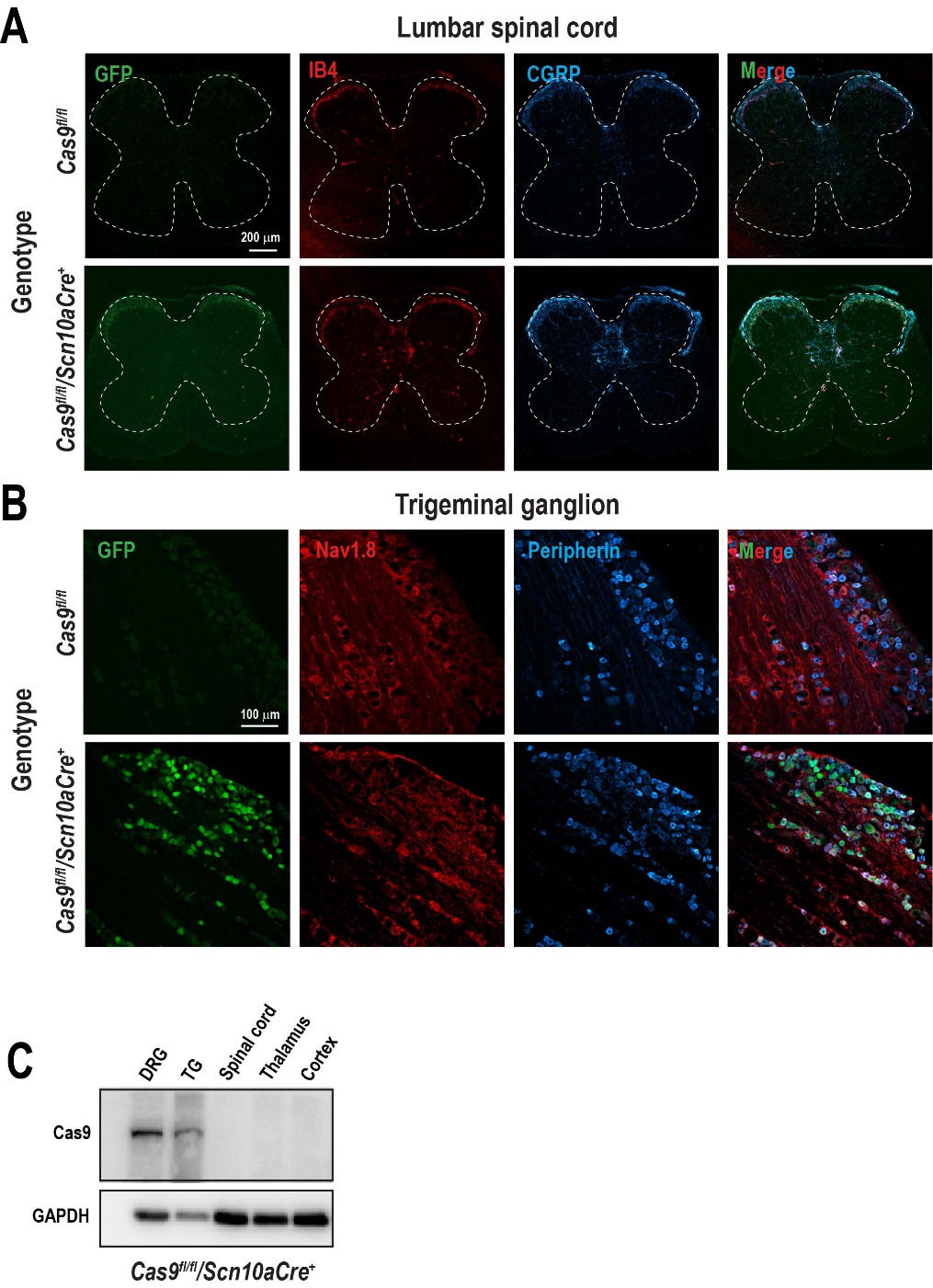

**Supplementary Figure 1.** GFP expression in the (**A**) lumbar spinal cord and (**B**) trigeminal ganglion. Tissues were extracted from *Cas9^fl/fl^/Scn910aCre^+^* mice, where Cas9 and GFP expression are induced in a Cre recombinase-dependent manner. No GFP expression is observed in the absence of Cre recombinase (*Cas9^fl/fl^*). Images show colocalization of GFP with IB4 and CGRP in the laminae I and II of the dorsal horn of the spinal cord. In the trigeminal ganglion, GFP was colabeled with Nav1.8 and peripherin. Images are representative of three replicates. Scale bar: 100 µm. (**C**) Cas9 protein expression was determined by immunodetection in DRG, trigeminal ganglion (TG), lumbar spinal cord, thalamus and cerebral cortex from *Cas9^fl/fl^/Scn910aCre^+^* mice. Cas9 expression was restricted to sensory neurons.

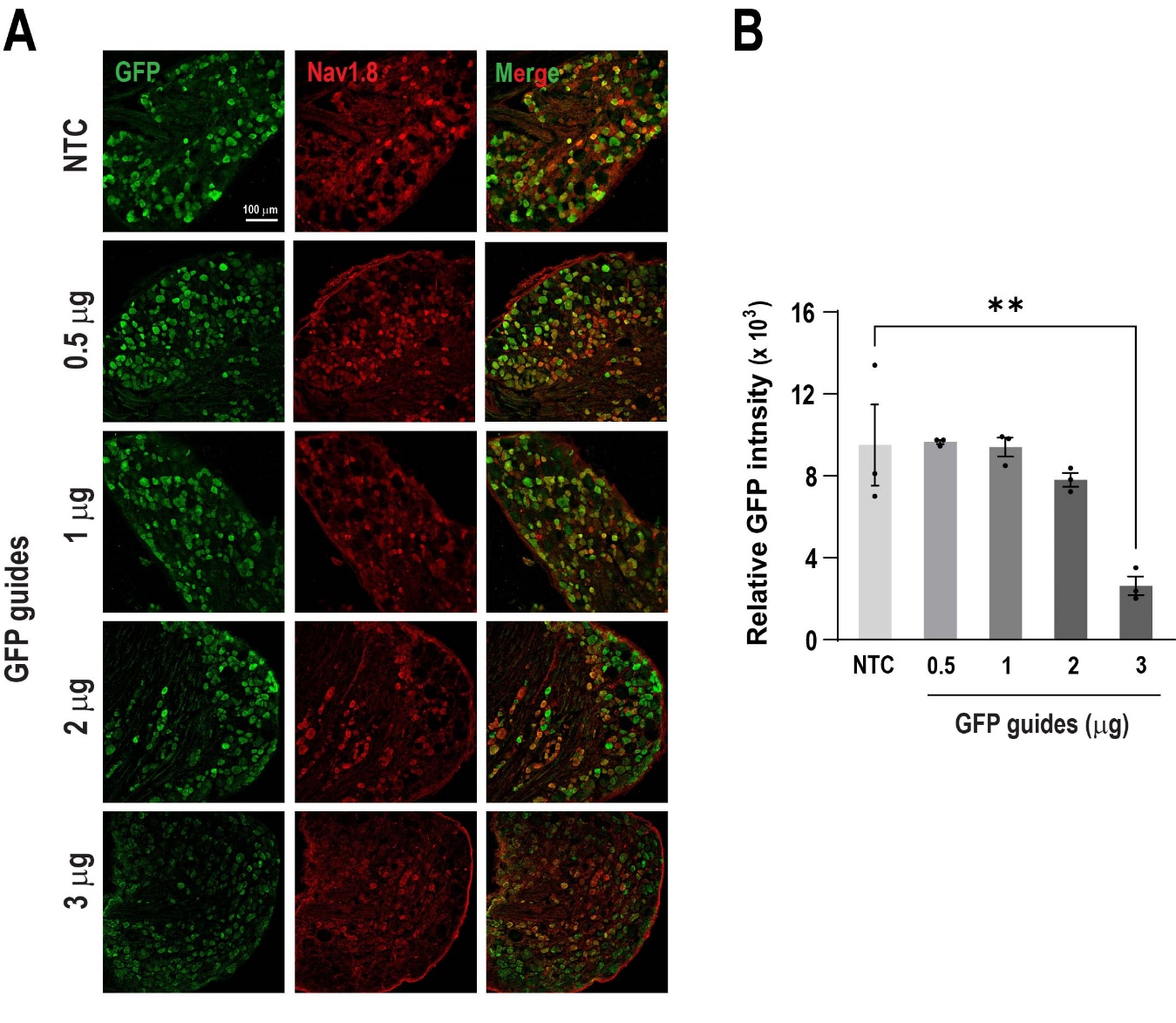

**Supplementary Figure 2.** Optimization of guide RNA dosage. (**A**) Representative images of immunohistochemical staining of DRGs following a single injection of increasing doses (0.5, 1, 2 and 3 µg) of GFP guides. Scale bar: 100 µm. (**B**) Quantitative analysis of relative GFP intensity normalized to Nav1.8 signal. The data show a significant reduction in GFP expression at the 3 µg dose compared to the NTC group. Bars represent the mean ± SEM of relative intensity values from n = 3 mice per group. ***P* < 0.01 was determined by one-way ANOVA followed by Dunnett’s test.

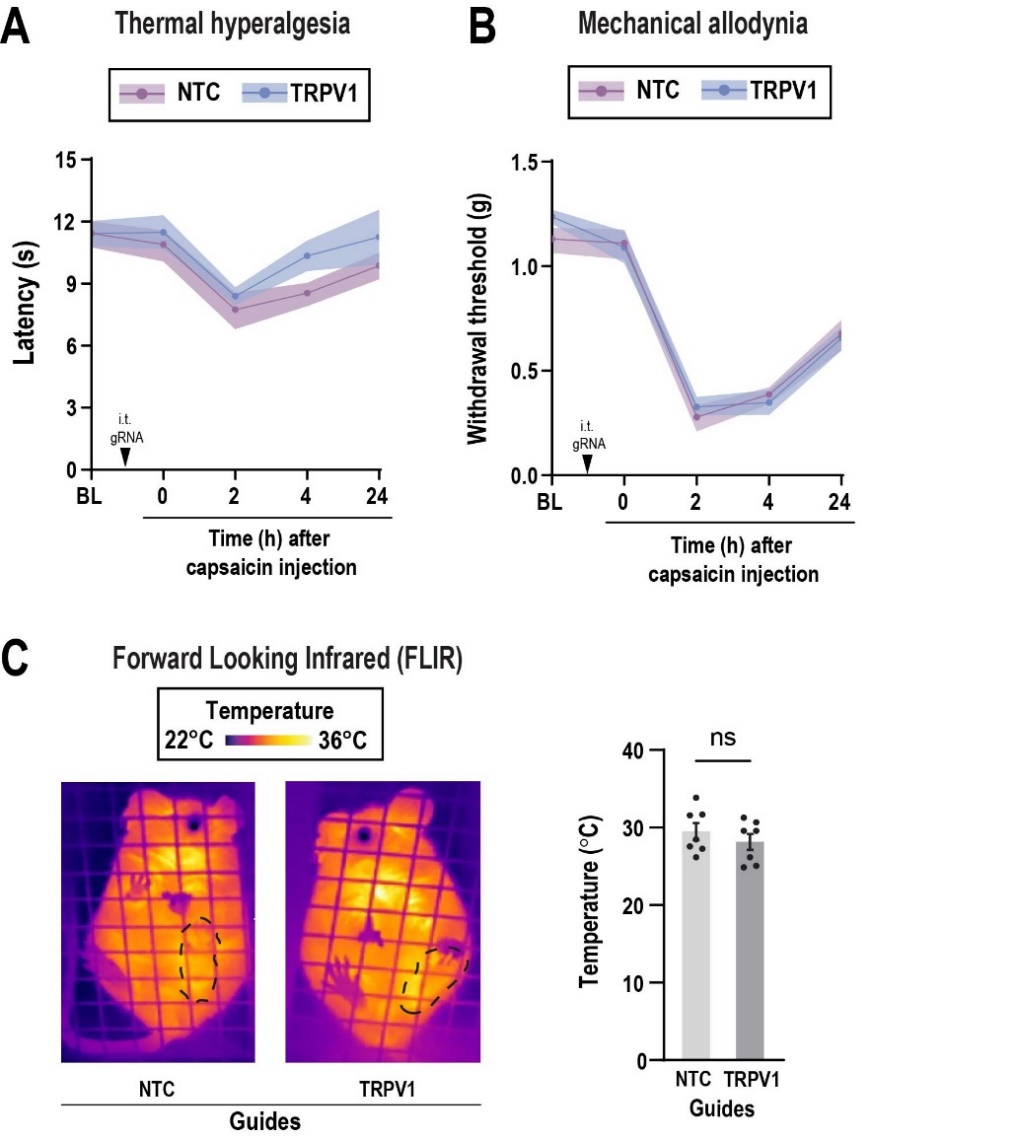

**Supplementary Figure 3.** A single injection of TRPV1 RNA guides is not sufficient to prevent capsaicin-induced hypersensitivity in Cas9 knock-in mice. Mice received one intrathecal injection of TRPV1 or NTC guides (3 μg). One week after the injection, behavioral tests were conducted. (**A**) Thermal hyperalgesia and (**B**) mechanical allodynia were evaluated before intrathecal injection of TRPV1 or NTC guides, as baseline (BL), and one week after guide delivery at 0, 2, 4, and 24 hours after capsaicin injection. Lines represent the mean ± shaded area indicating SEM of latency (s) or withdrawal threshold (g) from n = 7 mice per group. No significant difference was found following two-way ANOVA. Arrow indicates intrathecal injection of RNA guides (i.t. gRNA). (**C**) Representative images of the right paw injected with capsaicin (dotted line) from TRPV1 or NTC guide-treated mice. Forward looking infrared (FLIR) imaging was used to measure paw temperature 1 h after capsaicin injection. Bars represent the mean ± SEM of temperature (°C) from n = 7 mice per group. No significant difference (ns) was detected by an unpaired t-test.

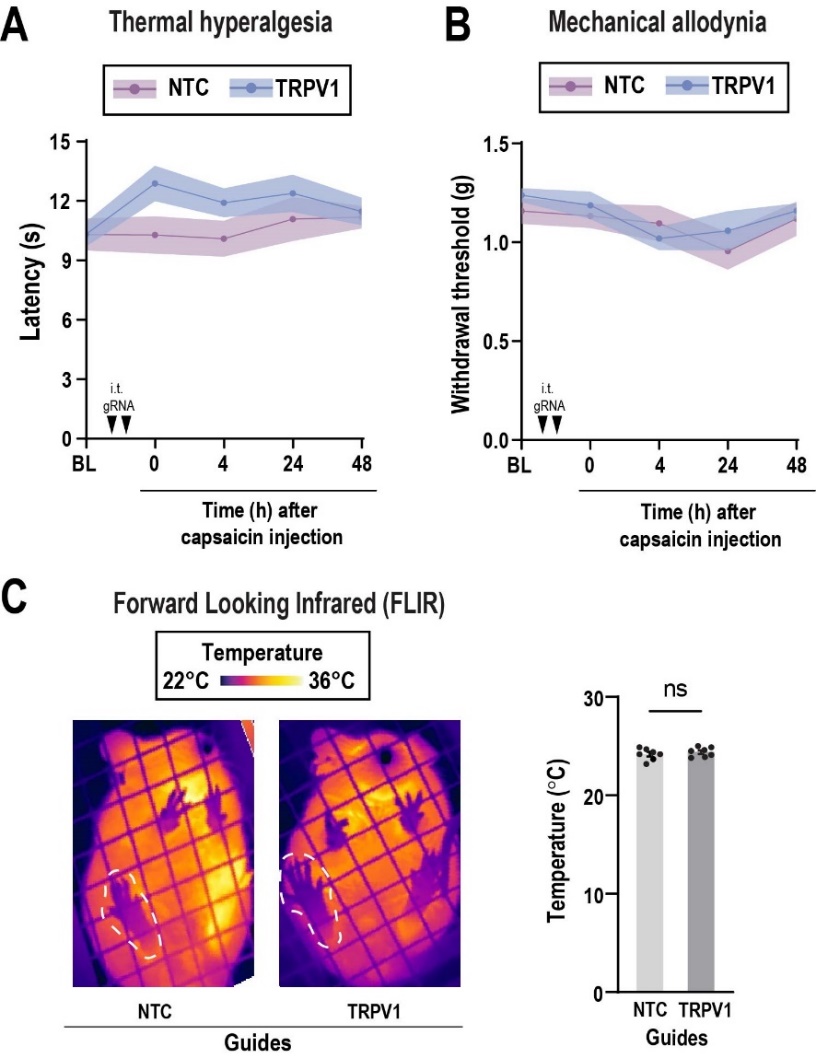

**Supplementary Figure 4.** Capsaicin does not evoke hypersensitivity and inflammation in the contralateral paw. Mice received two intrathecal injections of TRPV1 or NTC guides (3 μg per injection) over a period of 2 weeks. One week after the last intrathecal injection, 5 μg of capsaicin was intraplantarly injected and behavioral tests were conducted on the contralateral paw. (**A**) Thermal hyperalgesia and (**B**) mechanical allodynia were evaluated before intrathecal injection of TRPV1 or NTC guides, as baseline (BL), and one week after guide delivery at 0, 4, 24 and 48 hours after capsaicin injection. Lines represent the mean ± a shaded area indicating SEM of latency (s) or withdrawal threshold (g) from n = 6-7 mice per group. No significant difference was found following two-way ANOVA between treatments and time points. Arrows indicate intrathecal injections of RNA guides (i.t. gRNA). (**C**) Representative images of the right contralateral paw injected with capsaicin (dotted line) from TRPV1 or NTC guide-treated mice. Forward looking infrared (FLIR) imaging was used to measure paw temperature 1 h after capsaicin injection. Bars represent the mean ± SEM of temperature (°C) from n = 7 mice per group. No significant difference (ns) was detected by an unpaired t-test.
